# Individual *Staphylococcus aureus* SauUSI restriction endonuclease motors fragment methylated DNA

**DOI:** 10.1101/2025.10.16.682607

**Authors:** S.J. Shaw, A. Hughes-Games, O.E. Torres Montaguth, S. McDowall, F.M. Diffin, D. Dunn, S.J. Cross, M.D. Szczelkun

**Affiliations:** DNA-Protein Interactions Unit, School of Biochemistry, University of Bristol, Bristol, UK

## Abstract

SauUSI is a dimeric, ATP-dependent Type IV restriction enzyme that protects *Staphylococcus aureus* by cleaving non-self DNA containing 5-methylcytosine or 5-hydroxymethylcytosine. Using biophysical, single-molecule, and nanopore sequencing methods, we show that 5-methylcytosine recognition first induces ATP-driven unidirectional translocation by one helicase-like subunit, displacing its target recognition domain (TRD). The partner TRD can then bind the liberated modified site, stabilizing a growing DNA loop. On symmetrically methylated DNA, both subunits engage in bidirectional loop translocation. Cleavage is triggered by binding a distal methylated site, by preferred DNA sequences, or upon reaching a DNA end. Interactions with other SauUSI dimers or protein roadblocks affect cleavage site distributions but are not required for nuclease activation. While the first cleavages principally generate blunt-ends or one-nucleotide 3′ overhangs, multiple binding-translocation cycles by individual enzymes ultimately shred the modified non-self DNA, neutralizing its threat.

## INTRODUCTION

Bacteria and archaea deploy a broad arsenal of defense mechanisms to fend off invading bacteriophages (phages), plasmids, integrative elements, and other environmental genetic material^1–5^. To evade these defenses—especially Restriction-Modification (RM) and CRISPR-Cas—phages and other mobile genetic elements evolved a variety of chemical modifications to mask recognition sites and interfere with nuclease activity^6–13^: including base modifications such as N6-methyladenine, 5-methylcytosine (5mC), 5-hydroxymethylcytosine (5hmC), 5-glucosylhydroxymethylcytosine, atypical bases like uracil, or DNA backbone changes such as phosphorothioation. Under selective pressure, additional microbial defenses then evolved to recognize such modifications and mount a response^5,14–16^. Here we explored how the Type IV restriction endonuclease SauUSI can efficiently trigger multiple breaks in DNA following 5mC recognition.

SauUSI was originally identified in clinical *Staphylococcus aureus* strains as a major barrier to horizontal gene transfer (HGT) of methylated plasmids and to infection by methylated phages^17–19^. Restriction is thus likely to influence acquisition of virulence, pathogenicity and antimicrobial resistance genes on prophages, plasmids, etc, ^20,21^ and SauUSI homologues are found across Gram-positive and Gram-negative bacteria, and some archaea^22^. Unlike classical restriction enzymes that recognize specific unmodified nucleotide sequences^5^, SauUSI recognizes 5mC or 5-hmC within a degenerate sequence (5ʹ-SXNGS-3ʹ, where S = G or C and X is the modified base, defined using a finite set of target sequences) ^18^. As such, it is classified as part of the Type IV restriction endonucleases that only recognize and cleave modified DNA^5,14,23^. *In vitro* assays demonstrated that SauUSI cleaves DNA using ATP (or dATP) hydrolysis in the presence of one or more methylated sites^18,22^. The main dsDNA breaks were reported as mapping 5–20 bp away from the targets and multiple targets resulted in more efficient cleavage and DNA fragmentation (“shredding”) ^22^.

SauUSI consists of an N-terminal phospholipase-D (PLD)-family nuclease, dual Superfamily 2 (SF2) helicase-like RecA domains, a coupling domain, and a C-terminal Target Recognition Domain (TRD) with a SET and RING-associated (SRA) fold that recognizes modified bases (**Fig. 1A**). Structural studies showed two monomers dimerize via the PLD domains to form a single nuclease active site^22^, as seen in the PLD-based Type II restriction enzyme BfiI^24–26^. The SauUSI TRD likely flips the modified base into a recognition pocket, triggering ATP hydrolysis and dsDNA translocation by the helicase-like ATPase (rather than unwinding) ^18,22^, though the polarity of motion remains unclear. Cleavage was proposed to occur when translocating motors “pile up” against at a second site, each motor generating random breaks, ultimately shredding the DNA^22^. However, the need for multiple enzymes remains puzzling as a single PLD dimer should suffice to cut both DNA strands in a sequential mechanism^24,25,27^. It is more typically thought that pairs of translocating restriction enzymes interact to dimerize and thus activate nuclease domains^28–30^.

**Fig. 1.**
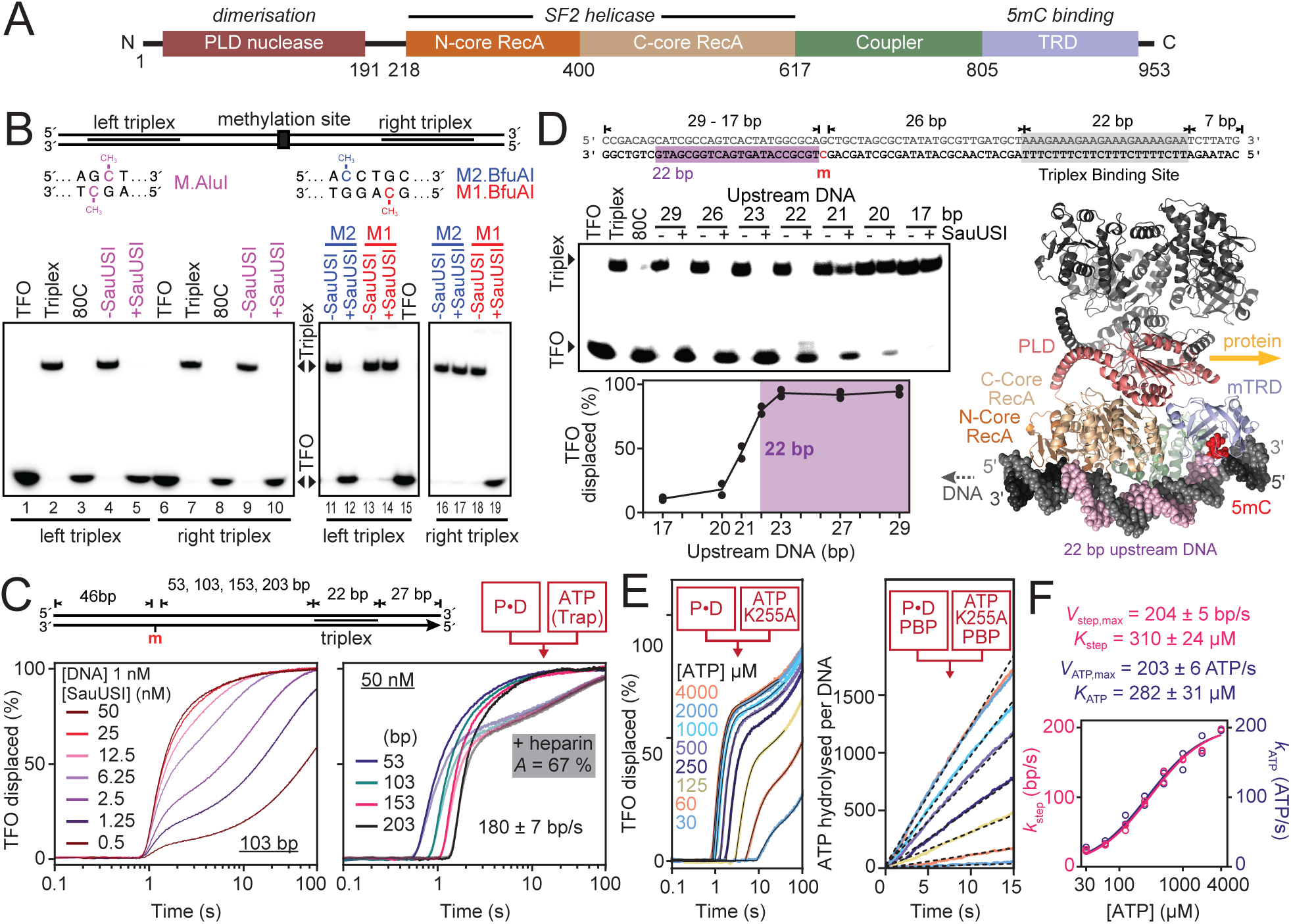
Methylated DNA strand dependent translocation. **A**, Primary domain structure of SauUSI. **B**, Translocation directionality from displacement of the triplex forming oligonucleotides (TFO) using triplex gel electrophoresis assays with dimethylated DNA (M.AluI) or strand-specific hemimethylated DNA (M1/M2.BfuAI). **C**, Time-dependent triplex displacement from linear DNA with different hemimethylation – triplex distances measured using a stopped flow assay. Samples were mixed as indicated (cartoon) with Protein (P), DNA (D), ATP and heparin (Trap, where included). Rate determined in **Supp. Fig. X**. **D**, Role of upstream DNA in supporting translocation using a triplex gel electrophoresis assay with linear DNAs with different length of DNA upstream of a hemimethylated site. Data points are the average of 2 repeats with a line joining the points. (*inset*) AlphaFold3 model of a SauUSI dimer with one subunit binding a 41 bp hemimethylated DNA. **E**, Effect of ATP concentration on initiation measured using stopped flow triplex displacement (*left panel*) or on ATP hydrolysis rates measured using the phosphate binding protein (PBP, *right panel*). Triplex assays were fitted to a plateau followed by a two-phase association (black line), with the amplitude of the fast phase giving the proportion of active translocating motors in the presence of excess K255A trap. The ATPase data was corrected for background ATPase activity and fitted by linear regression (black dotted line) to give rates that were adjusted by the active motor fraction (**Extended data Fig. 2B,C**). **F**, Comparison of the ATP dependence of the translocation rate (*k*_step_) and ATPase rate (*k*_ATP_). Lines are separate hyperbolic fits to the data (n = 2). Quoted parameters are the mean and standard errors.

To better understand SauUSI defense, we used biochemical, single-molecule and nanopore sequencing assays. Our data show unidirectional 3ʹ–5ʹ translocation along a methylated strand, that can also form expanding DNA loops, while methylation on both strands can support bidirectional loop translocation. Cleavage occurs both proximal and distal from 5mC sites, driven by local sequence rather than dimer interactions or roadblock collisions. Repeated cycles of 5mC recognition and motor activity generate multiple random dsDNA breaks, characteristic of shredding, explaining the potent restriction of methylated DNA seen in *Staphylococcus aureus* cells.

## RESULTS & DISCUSSION

### SauUSI is a directional dsDNA translocase that initiates upstream of 5-methylcytosine

To confirm translocation directionality, we monitored triplex displacement^31^, using triplex binding sites flanking methylated targets that were either dimethylated by AluI methyltransferase (M.AluI) or hemimethylated by the strand-specific methyltransferases M1.BfuAI (bottom strand) or M2.BfuAI (top strand) (**Fig. 1B**). On dimethylated DNA, SauUSI displaced both left and right triplexes (lanes 5 and 10). On hemimethylated DNA, displacement was methylated strand-dependent: with M2.BfuAI (top strand), only the left triplex was displaced (lanes 12 vs. 17); while, with M1.BfuAI (bottom strand), only the right triplex was displaced (lanes 14 vs. 19). Translocation is thus directional, proceeding 3ʹ–5ʹ along the methylated strand. When both strands are methylated, translocation in both directions is possible, as explained below.

To measure translocation rates, we used fluorescent triplexes in a stopped-flow fluorimeter^32^, with SauUSI pre-bound to DNA containing varying distances between a hemimethylated site and a triplex binding site, and initiated reactions by rapid ATP mixing (**Fig. 1C**). At different SauUSI concentrations (left panel), a consistent lag-phase preceded triplex displacement, indicating stepwise translocation. However, saturation required excess enzyme, suggesting that DNA-binding limits initiation. With excess enzyme, the lag time increased linearly with site-triplex distance, yielding a translocation rate of 180 ± 7 bp/s at 25 °C (right panel, **Extended data Fig. 1**). A trap could be included with ATP to prevent dissociated enzymes rebinding - either heparin or a methylated oligoduplex (M-Oligo) to block the TRD(s) or an ATPase-deficient SauUSI mutant (K255A) ^22^ to bind the released 5mC (**Supp. Fig. 1**). The displacement amplitude with trap was uniform with distance (e.g. **Fig. 1C**, right), implying ~30% of enzymes dissociated before/during initiation, while dissociation during translocation was negligible over these distances (the translocation rate was >1000-fold faster).

When the triplex profiles were fitted to translocation models (**Supp. Fig. 2**) ^32,33^, the number of 1 bp steps needed to produce consistent translocation rate estimates was greater than expected from motor initiation immediately downstream of the 5mC. An explanation is that the SauUSI motor binds upstream, increasing the initiation site-to-triplex distance. We tested this using DNA with fixed 5mC– triplex spacing but varied upstream DNA lengths (**Fig. 1D**). Efficient displacement required ≥22 bp upstream. Using AlphaFold3 (ref^34^) to produce a DNA-bound structural model supports upstream initiation, showing ~22 bp between a flipped 5mC and the ATPase-DNA contact point (**Fig. 1D**). Upstream initiation gives more consistent estimated translocation rates (**Supp. Fig. 2**). With 3ʹ–5ʹ translocation along the methylated strand, the motor will move toward the TRD (**Fig. 1D**). Given the geometry and expectation of interactions with both strands, the motor likely cannot bypass the TRD without displacing it.

### A single helicase-like ATPase motor is active during unidirectional dsDNA translocation

To demonstrate that only one motor in the dimer is active on hemimethylated DNA, we measured translocation and ATPase rates at sub-saturating ATP concentrations (**Fig. 1E**, **Extended data Figs. 1 & 2**), using excess K255A to prevent multiple events reinitiating. As ATP decreased, dissociation increased (**Fig. 1E, left**), and this relationship was used to correct the ATP hydrolysis rates (**Fig. 1E, right**; **Extended data Fig. 2C**). Both translocation and ATPase rates showed similar hyperbolic ATP dependence (**Fig. 1F**), yielding a coupling of ~1 ATP per bp (**Extended data Fig. 2C**), consistent with a single subunit driving unidirectional movement.

**Fig. 2.**
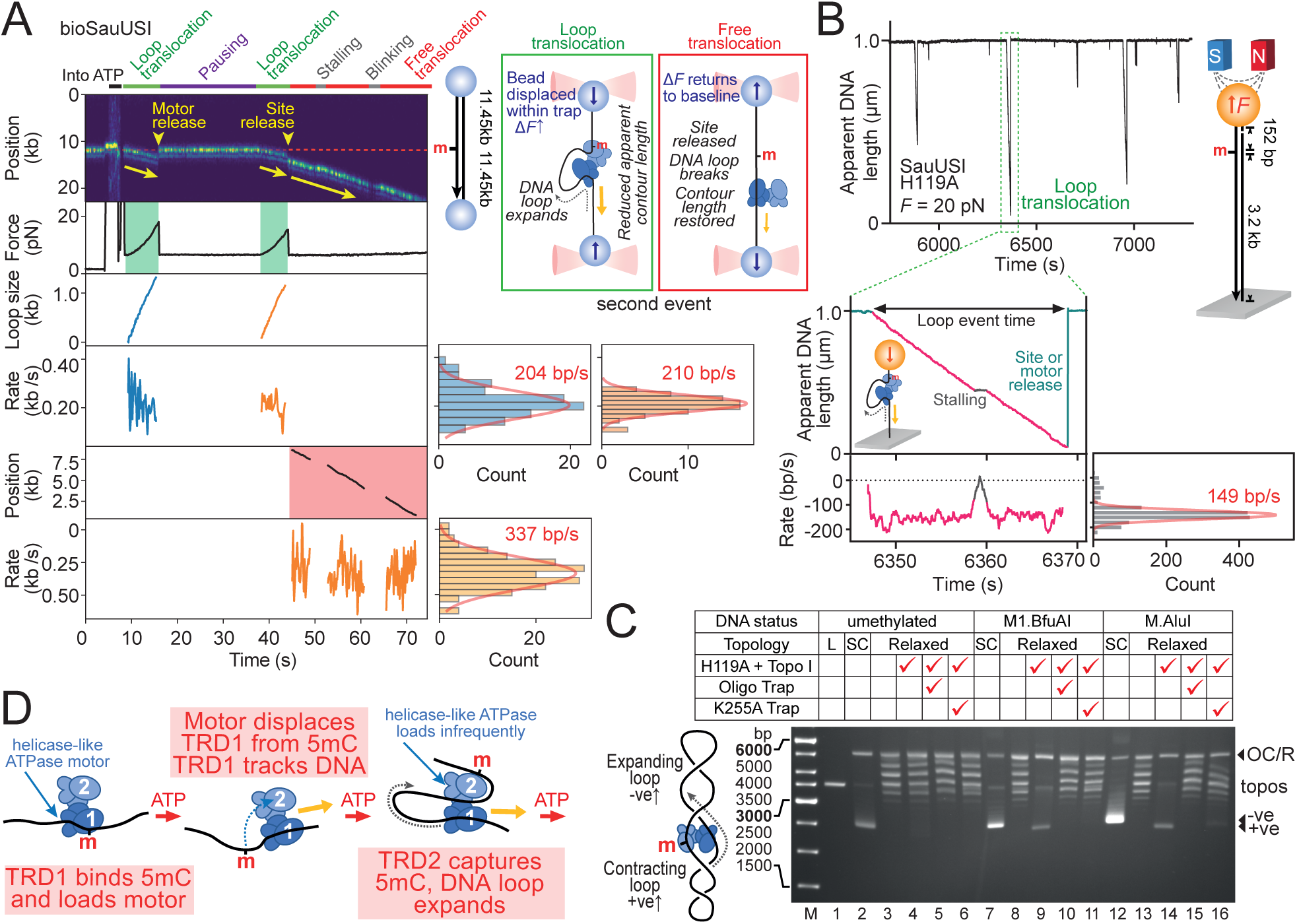
Translocation on hemimethylated DNA can result in unidirectional loop translocation. **A**, Representative kymograph at 1.5 pN showing a hemimethylated DNA-bead tether with labelled enzyme (655 nm quantum dot-streptavidin-biotin-WT SauUSI) prebound. After moving into the ATP channel, two loop translocation events (inset cartoon) are observed corresponding with increases in force. Free translocation (inset cartoon) indicates motion without a force change. Quantum dot blinking produces occasional trace gaps. Instantaneous rates for loop or free translocation events were binned to give average rates from Gaussian fits (bin widths were 31, 19, and 47 bp/s for derivation of rates 204, 210, and 337 bp/s, respectively). For details of single molecule parameters, see **Extended data Fig. 1**. **B**, Representative time trace from MT measurement of apparent DNA length at 20 pN showing DNA looping events by SauUSI H119A. Instantaneous rates for loop translocation events were binned and average rates calculated from multiple events. For details of single molecule parameters, see **Extended data Fig. 1**. **C**, Representative agarose gel electrophoresis of reactions using relaxed circular DNA (R) and controls in the presence of *E. coli* DNA topoisomerase I to provide evidence of loop translocation (cartoon). Reactions were initiated by adding ATP or ATP plus trap (M-Oligo or SauUSI-K255A). +ve/-ve are positions of positive and negative supercoiled DNA, OC is nicked DNA (open circle) and topos indicates topoisomers **D**, Model for how loops are formed during initiation of translocation. See text for full details. Quoted parameters are the mean and standard deviation.

### SauUSI can initiate unidirectional loop translocation on hemimethylated DNA

To further investigate translocation, we used a dual optical trap and fluorescence single-molecule setup (Lumicks C-trap) with ~23 kb biotin end-labelled hemimethylated DNA tethered between streptavidin-coated beads (**Fig. 2A**). SauUSI–quantum dot complexes were loaded onto DNA and transferred to ATP. Unidirectional translocation initiation was observed but in 46% of cases it coincided with increased DNA tension, indicating formation of an expanding DNA loop (**Fig. 2A**, **Extended data Fig. 3A-C**). In **Fig. 2A**, the first looping event ends with motor release returning SaUSI to the site (13% of all events), while the second event continues from a distant site after loop collapse, consistent with 5mC release followed by “free translocation” (87% of all events). Free translocation often ran thousands of bp to a DNA end, supporting processive translocation inferred from the triplex assay.

To rule out that quantum dot labelling facilitates looping, we used an unlabeled nuclease mutant (H119A) ^22^ with a magnetic tweezers (MT) microscope and a 3.2 kb DNA with a single hemimethylated site (**Fig. 2B**). DNA shortening events consistent with directional loop translocation were observed at a range of DNA stretching forces (**Extended data Fig. 3D-G**)-free translocation could not be measured in this set-up.

Translocation in 1 bp steps would reduce twist in an expanding DNA loop and increase twist ahead, generating separate supercoiled domains on circular DNA (**Fig. 2C**) ^35–37^. Including *Escherichia coli* Topoisomerase I to remove negative supercoils detected loop formation by conversion of relaxed to supercoiled DNA. Supercoiling was only observed with hemi- or dimethylated sites (lanes 4 versus 9 and 14). Adding K255A or M-Oligo traps reduced or eliminated supercoiling (lanes 10, 11, 15, 16), indicating that loop formation depends on both the TRD(s) and a liberated 5mC.

In a model for hemimethylated DNA loop translocation (**Fig. 2D**), following TRD1-5mC binding, the upstream motor initiates 3ʹ–5ʹ translocation, displacing TRD1. In half of the events, free translocation continues. Cleavage data, below, is consistent with TRD1 tracking the DNA once remodeled, blocking site recapture after initiation. Instead, TRD2 binds the released 5mC in the other half of events forming a loop (**Fig. 2B**, **Extended data Fig. 3D-F**). The traps used in **Fig. 2C** prevent 5mC capture by TRD2—either via blocking entry to TRD2 (M-Oligo) or by binding and shielding the released 5mC (K255A). Since looping occurs at forces that resist DNA bending, the second subunit must have conformational flexibility, allowing DNA binding (not shown in the cartoon). Although events consistent with co-directional movement of two motors were observed, suggesting the second motor can initiate (e.g. **Supp. Fig. 3**), these were rare (n = 4) and the loop geometry must normally prevent initiation.

### SauUSI can initiate bidirectional loop translocation on dimethylated DNA

Triplex displacement on both sides of the M.AluI site (**Fig. 1B**) could result from random unidirectional translocation or simultaneous bidirectional activity. Both events were supported by single-molecule assays with DNA containing a single dimethylated site (**Fig. 3**).

**Fig. 3.**
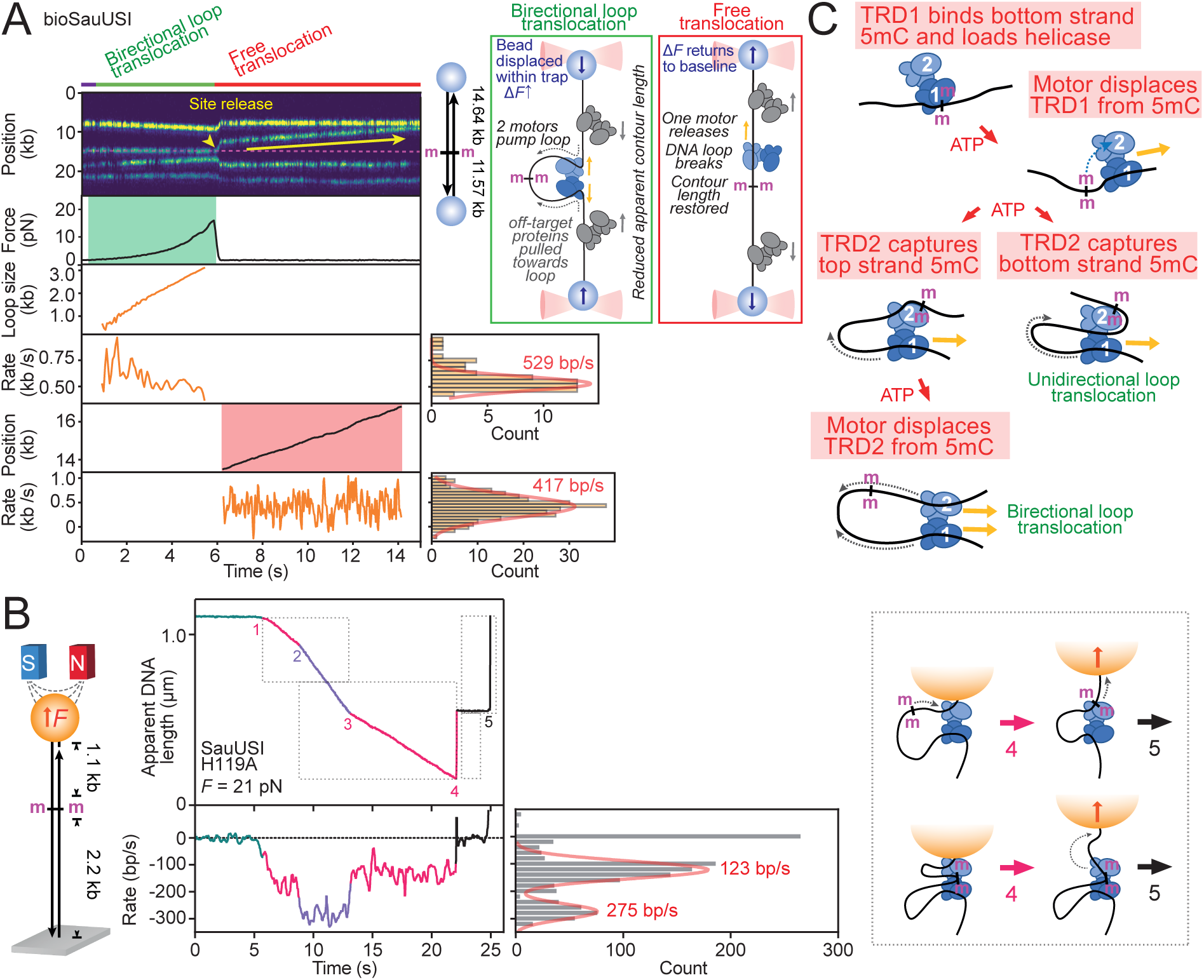
Translocation on dimethylated DNA can result in bidirectional loop translocation. **A**, Representative kymograph at 1.5 pN showing a dimethylated DNA-bead tether with labelled enzymes (655 nm quantum dot-streptavidin-biotin-WT SauUSI) prebound at the 5mC and at off-target sites. Following a brief delay after moving into the ATP channel, a bidirectional loop translocation event (inset cartoon) is observed corresponding with an increase in force. Free translocation (inset cartoon) indicates motion without a force change. Quantum dot blinking produces occasional trace gaps. Instantaneous rates for loop or free translocation events were binned to give average rates from Gaussian fits (bin widths were 40 and 64 bp/s for derivation of rates 529 and 417 bp/s, respectively). For details of single molecule parameters, see **Extended data Fig. 1**. **B**, Representative time trace from MT measurement of apparent DNA length at 21 pN showing DNA looping events by SauUSI H119A on dimethylated DNA. Instantaneous rates for loop translocation events were binned and average rates calculated from multiple events. For details of single molecule parameters, see **Extended data Fig. 1**. Dotted boxes represent events during which one or two motors are active/bound (see main text). (*inset cartoon*) Two alternative explanations for event 4 (see main text). For details of single molecule parameters, see **Extended data Fig. 1**. **C**, Model for how initiation of translocation on dimethylated DNA may result in either uni- or bidirectional translocation. See text for full details.

Using a dimethylated DNA, four labelled SauUSI are visible in the C-trap example in **Fig. 3A**— one 5mC-bound and three non-specific. An increase in force indicates looping, but, distinct to **Fig. 2A**, the target-bound enzyme appears stationary while the others are pulled inward, consistent with bidirectional translocation by two motors. Loop collapse leads to free translocation, “upwards” in **Fig. 2A**, with off-target enzymes returned to their static positions, suggesting release by the “downward” motor. It was, however, difficult to assign the dissociating domain in most events. Compared to hemimethylated DNA, there was a higher frequency of looping (73%) of which at least half were bidirectional (**Extended data Fig. 3B,C**).

In the MT example in **Fig. 3B**, an initial unidirectional event is followed by a faster phase, indicating activation of the second motor and bidirectional translocation. When one motor stalls at the bead, the rate drops back to that of a single motor. At ~22 seconds, the rapid increase in DNA height may reflect either (inset cartoon): (1) a single loop where one motor back-slides and rebinds 5mC; or, (2) dual-loop translocation where the TRD(s) bind the methylated site throughout, with one motor releasing. After a brief stall, the remaining loop collapses. Bidirectional looping was observed with similar frequency (21-25%) at DNA stretching forces of 9-21 pN (**Extended data Fig. 3H**).

A model for bidirectional loop translocation on dimethylated DNA is shown in **Fig. 3C**. After 5mC binding, the upstream motor initiates 3ʹ–5ʹ translocation, displacing TRD1 that remains DNA associated. Looping then occurs via TRD2 rebinding to either: (1) the bottom strand 5mC (same strand as the first motor), leading to unidirectional loop translocation as the second motor cannot load; or (2), the top strand 5mC, enabling second motor loading and bidirectional translocation.

### DNA loop translocation produces lowers motor rates than free translocation

Unidirectional loop translocation by wild-type SauUSI averaged 256 ± 54 bp/s in the C-trap (ambient temperature), while H119A averaged 132 ± 14 bp/s in the MT (25 °C) (**Extended data Fig. 3A,G**). H119A did not show slower rates in the C-trap, suggesting the difference may stem from local sample heating by C-trap lasers. Based on a Q10 temperature coefficient of 1.92 from triplex displacement data (**Supp. Fig. 4**), a C-trap flow cell temperature of ~35 °C would account for the rate difference. In both assays, forces up to 20 pN had no effect on translocation rates, consistent with other DNA motors that translocate with small step sizes (1 bp ≈ ~0.34 nm) ^38^. While force affected the lifetime of longer-lived loops in the MT (**Extended data Fig. 3E**), the trend was opposite to that expected from loop strain, instead likely reflecting experimental variation. Additionally, in the tweezers ~29-50% of hemimethylated DNA looping events were short-lived (<1.5 s) but were not observed using dimethylatyed DNA or in the C-trap, possibly reflecting unstable H119A-specific loops or early initiating free translocation events.

Bidirectional loop translocation by wild-type SauUSI averaged 548 ± 59 bp/s in the C-trap (ambient temperature), and 306 ±35 bp/s for H119A in the MT (25 °C)—slightly more than double the corresponding unidirectional rates, suggesting some allosteric activation rather than completely independent motors (**Extended data Fig. 3A,G,H**). A rate-force dependence could not be confidently determined.

Free translocation in the C-trap averaged 317 ± 58 bp/s for wild type and 363 ± 46 bp/s for H119A (**Extended data Fig. 3A**). These rates were significantly higher than unidirectional looped translocation rates, suggesting that loop geometry might slow motor activity. Free translocation may only occur on stretched DNA; at lower tension (or DNA “in solution”), the free TRD may rebind distant sites via DNA flexing.

### DNA translocation transports the nuclease to the site of cleavage

To investigate nuclease activation, we first analyzed hemimethylated DNAs. On supercoiled DNA with one bottom strand hemimethylation (named SR1B), SauUSI first linearized the ring (“primary cleavage”), followed by random fragmentation (“secondary cleavage”), regardless of enzyme concentration—only the rate varied (**Fig. 4A**). Equivalent cleavage patterns and rates were seen using linear DNAs, with products consistent with unidirectional downstream translocation and secondary shredding ending with products terminated near the 5mC (**Fig. 4B**). Cleavage also occurred on linear DNA with hairpin ends lacking free termini, indicating that activation is independent of 5ʹ phosphate or 3ʹhydroxyl groups (**Supp. Fig. 5**).

**Fig. 4.**
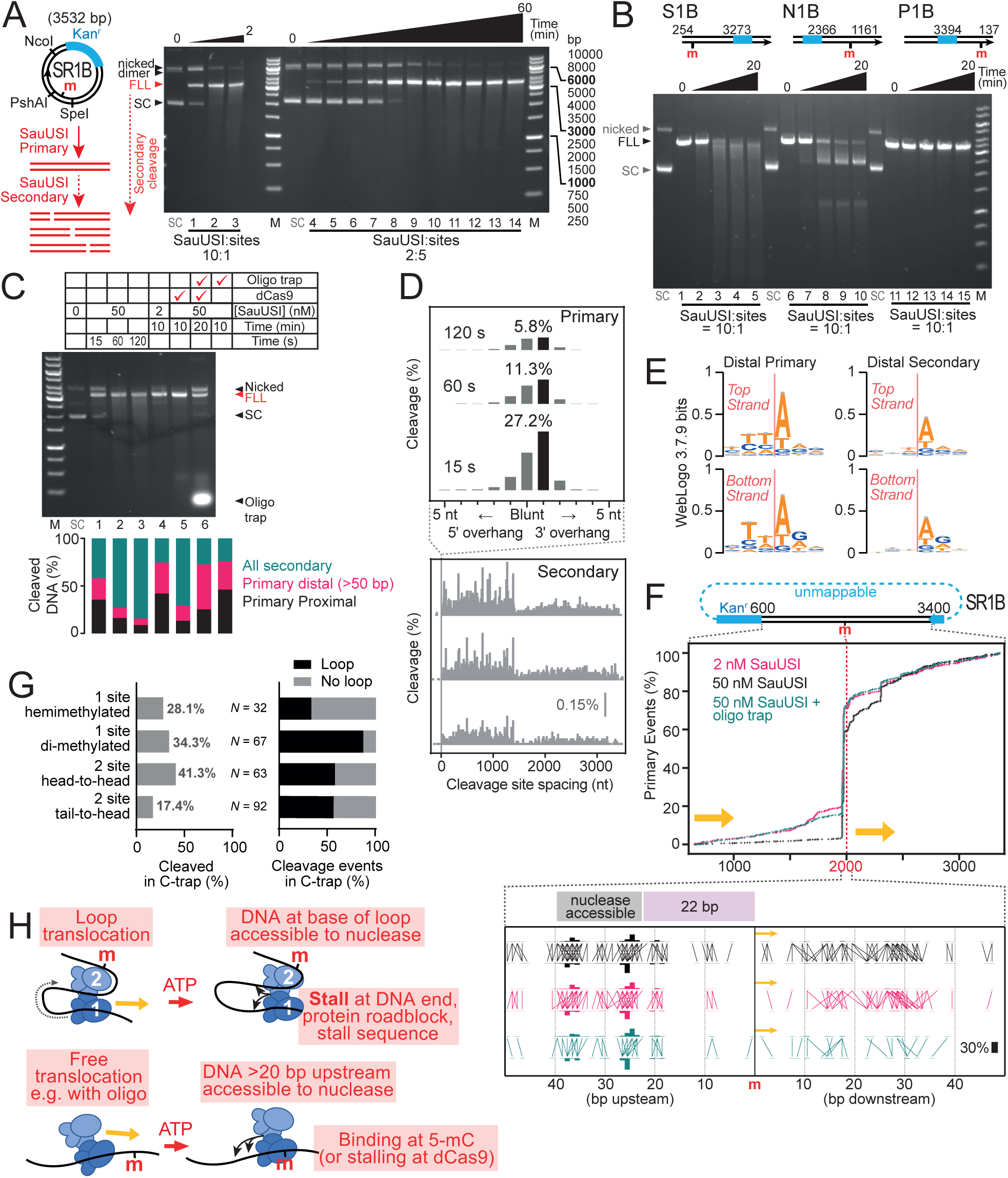
DNA cleavage by SauUSI is directed by translocation to both target site proximal and distal locations. All cleavage reactions were initiated by ATP. **A**, Time-course of cleavage of hemimethylated SR1B by sub-stoichiometric or excess SauUSI, measured using agarose gel electrophoresis. Cartoon shows SR1B map and products from the first (primary) cleavage that cuts the supercoiled (SC) DNA to produce full length linear (FLL) DNA via a nicked intermediate and subsequent (secondary) random cleavages that results in a smear of DNA fragments. Time points were 0.25, 1, 2 min (10:1 SauUSI:sites) and 0.25, 1, 2, 5, 10, 20, 30, 45 & 60 min (10:1 SauUSI:sites). **B**, Time-course of cleavage of hemimethylated DNA linearised with SpeI (S1B), NcoI (N1B) or PshAI (P1B), measured using agarose gel electrophoresis. Cleavage limit products tend towards fragments ending at the 5mC site. Time points were 0, 0.25, 2, 5 and 20 min. **C**, Agarose gel of reactions used for ENDO-Pore and event types quantified from the ENDO-Pore data (see main text for definitions). **D**, Distribution of cleavage site spacings from ENDO-Pore for the 15, 60 and 120 s reactions with excess (50 nM) SauUSI (1 nt bins). **E**, WebLogos from nucleotide frequencies 3 bp up- and downstream of top and bottom strand proximal and secondary cleavage; distal sites were analyzed as proximal cleavage is biased by 5mC binding (see panel F). **F**, Cumulative percentage of primary events relative to the 5mC positi on (m) on SR1B. Cleavage within the Kanamycin resistance cassette cannot be mapped by ENDO-Pore. Yellow arrow indicates polarity of unidirectional translocation. For the proximal region, a strand linkage plot (SLP) shows bars representing the percentage of phosphodiester cleavage at each top and bottom location and horizontal/vertical lines that indicate the ends generated (blunt or overhangs). **G**, Cleavage events types scored in the C-trap. **H**, Model to explain cleavage loci for looped and free-translocating species (see main text for full details).

To map cleavage products on supercoiled DNA, we used nanopore-based ENDO-Pore^39^ (**Fig. 4C**). Primary cleavage of hemimethylated circular DNA mainly produced blunt ends or 1-nt 3ʹ extensions (**Fig. 4D, Supp. Fig. 6**), similar to PLD-nuclease BfiI (Ref^40^). The extension variability here likely reflects repositioning of the PLD dimer during sequential strand cutting combined with local sequence preference (see below).

With either excess or sub-stoichiometric SauUSI, primary cleavage occurred throughout the mappable DNA but with sequence preference and favored loci (**Fig. 4E,F, Supp. Figs. 7 & 8**). We defined two primary cleavage loci: “proximal” (±50 bp from the 5mC) and “distal”. Distal cleavages were enriched at AT-rich sites, especially 5ʹ to dT or dA (**Fig. 4E, Supp Fig. 8A**), and occurred with lower frequency as distance from the 5mC increased (**Fig. 4F**), consistent with translocation directionality. High concentration produced more cleavage near the site compared to low, likely due to motor crowding (see below). Proximal cleavage was biased toward a 20 bp window ~22 bp upstream of the 5mC, with less frequent cuts downstream, refining previous observations^22^ (**Fig. 4F**). The upstream loci align approximately with the nuclease position relative to 5mC in the AlphaFold3 model (**Fig. 1D**). Similar cleavage patterns were observed with top strand hemimethylation (SR1T), although with different proximal and distal cleavage site preferences likely due to the opposite translocation polarity (**Supp. Fig. 9**)

DNA cleavage in the C-trap was comparatively infrequent but preceded by both looped or free translocating events (**Fig. 4G**, **Extended data Fig. 3C**). Stalling that did not result in DNA cleavage was regularly observed; DNA tension may be inhibitory.

Including M-Oligo to block looping (**Fig. 2C**) and reinitiation (**Supp. Fig. 1**) reduced overall cleavage but still produced primary and secondary events at levels equivalent to low SauUSI (**Figs. 4C,F**). Primary cleavage was distributed across distal loci but concentrated at proximal upstream sites (**Fig. 4F**), consistent with free translocation around the ring and 5mC reassociation. We propose that while M-Oligo can bind TRD2 and block loops forming (**Fig. 2C**), TRD1 stays DNA-associated during translocation, preventing *in trans* M-Oligo binding but allowing *in cis* 5mC recognition or stalling at other sequences.

A primary cleavage model has the nuclease triggered by a combination of local sequence preference and long-lived motor stalling (**Fig. 4H**). For looped events, the nuclease accesses sites at the loop base—both distal from and proximal to the 5mC. For free translocation, cleavage occurs only near the motor location. Upstream proximal cleavage (**Fig. 4F**) may result from the motor circling the DNA and the TRD that tracks the DNA ahead of the motor rebinding the 5mC, producing a stable state with efficient nuclease activity that supersedes any dinucleotide preference.

### Stalling against protein roadblocks biases cleavage loci but does not activate the nuclease

To test whether stalling at nucleoprotein roadblocks generally activates cleavage^22^, we tested site-specifically bound *Streptococcus pyogenes* dCas9 oriented with the PAM site facing the motor; some triplex displacement occurred when oriented oppositely (**Supp. Fig. 10**).

On linear and circular DNA with dCas9 bound 179 bp downstream from the 5mC, cleavage was mostly confined to the proximal region and up to the PAM (**Fig. 5**). Following loop translocation and stalling at dCas9, cleavage at the base of the loop would explain the elevated events in the downstream and upstream proximal regions and ~22 bp from the PAM (**Fig. 5B**). Assuming a low frequency of bypass, upstream proximal events may have resulted from motors circumnavigating the ring and rebinding the 5mC.

**Fig. 5.**
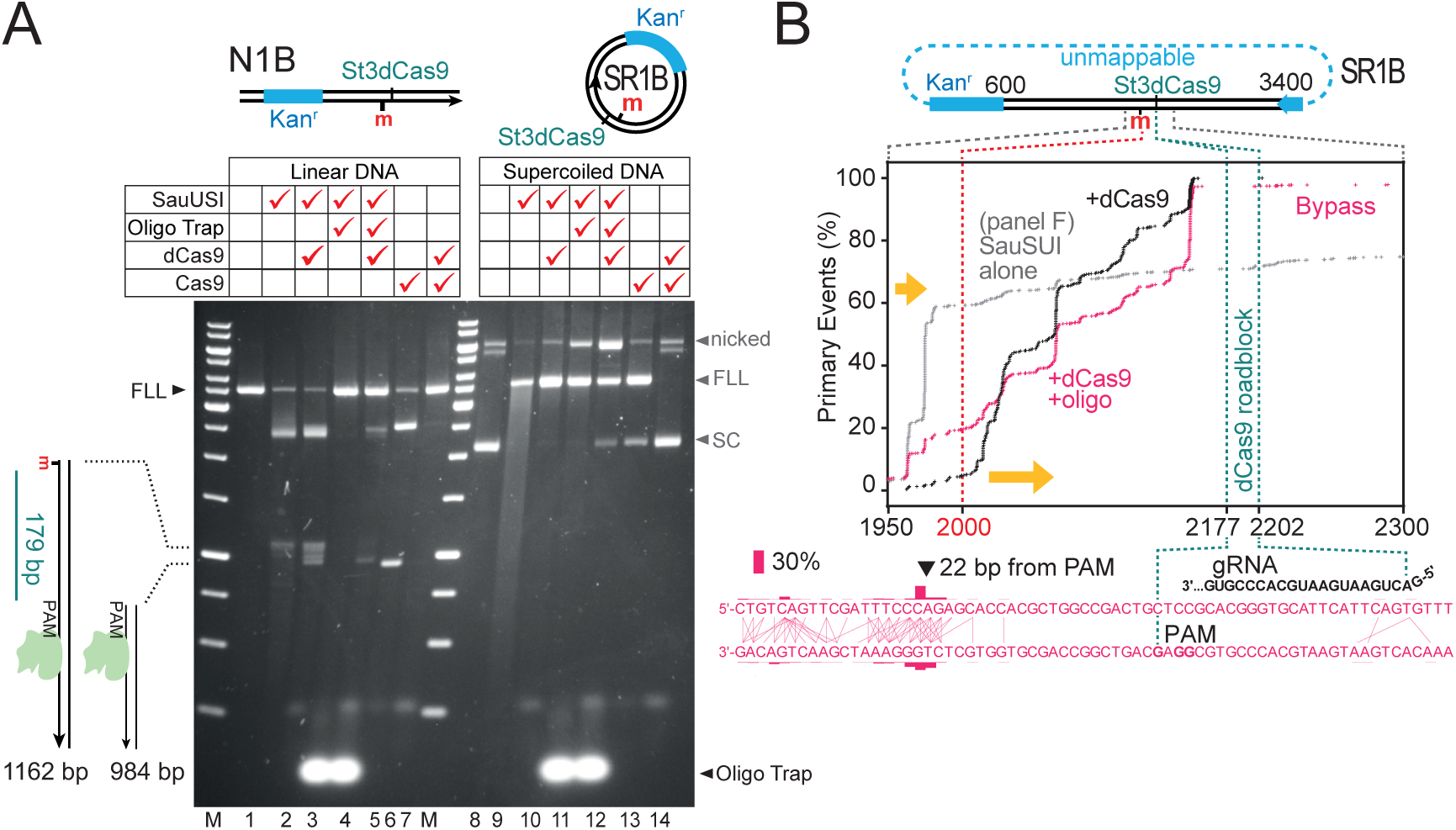
A dCas9 roadblock biases cleavage loci but does not activate the nuclease. **A**, Comparison of the effect of a dCas9 roadblock and M-Oligo trap on cleavage of SR1B and N1B. The Cas9 site was oriented with the PAM site facing the translocating SauUSI (**Supplementary Fig. 10**) DNA was incubated with excess SauUSI and with dCas9 as indicated, and 5-(2/4/9/11) or 10-(3/5-7/10/12-14) min reactions initiated with ATP, with trap where indicated. Binding of dCas9 was checked by its ability to block cleavage by Cas9. DNA was separated by agarose gel electrophoresis. The positions of the Cas9-dependent products from linear DNA cleavage are indicated. Gel markers as in **Fig. 2A**. **B**, Cumulative percentage of primary events relative to the 5mC position (m) on SR1B for reactions with dCas9 and with cCas9 plus M-Oligo. The proximal region to the dCas9 site is shown where most primary cleavage occurs. Yellow arrow indicates polarity of unidirectional translocation. For Cas9 site and region upstream of the PAM, a strand linkage plot (SLP) shows bars representing the percentage of phosphodiester cleavage at each top and bottom location and horizontal/vertical lines that indicate the ends generated (blunt or overhangs).

On circular DNA with dCas9 and M-Oligo trap, cleavage was reduced and confined to loci between the 5mC and dCas9 (**Fig. 5**). Fewer cuts occurred downstream of the 5mC (that we suggest are typical of looped events that are blocked here), while more appeared ~22 bp from the PAM, consistent with stalling of free translocating species and the structural model (**Fig. 1D**).

On linear DNA, M-Oligo blocked detectable cleavage (**Fig. 5A**), suggesting that without looping, translocation that terminates at DNA ends does not produce measurable cleavage, and stalling before the ends is too infrequent and/or distributed to observe using gel electrophoresis. Adding dCas9 then produced some detectable cleavage near the PAM, consistent with stalling of a free translocating motor (**Fig. 5A**). However, alongside the observation of reduced cleavage of circular DNA (above), a roadblock determines where cleavage can occur but does not significantly activate cleavage. Similarly, increased motor occupancy at elevated concentrations biases cleavage loci due to crowding rather than activation (**Fig**. **4F**). The concentration dependence of cleavage rates (**Fig. 4A**) more likely reflects second order limits on initiation rates (**Fig. 1C**, left panel).

On circular DNA with a single dimethylated site, low SauUSI cleaved efficiently within 5 minutes despite the reduced chance of collisions (**Extended data Fig. 4A**). Cleavage patterns on both sides of the 5mC were consistent with bidirectional translocation, with proportionally more distal cuts than on hemimethylated DNA (**Extended data Fig. 4**) but producing similar ends (**Supp. Fig. 7**). We interpret this as distal DNA being located at the base of bidirectionally translocated loops during cleavage (**Extended data Fig. 4C**), though unidirectional and free translocation events may also contribute.

In summary, we propose three modes of DNA cleavage activated by looped and non-looped species: (1) re-binding of a free 5mC site, producing efficient upstream cleavage; (2) stalling at a site with a preferred sequence for cleavage; (3) stalling at a DNA end. Crowding by multiple SauUSI motors or stalling at other roadblocks can determine where cleavage occurs through a corralling effect but is unnecessary for cleavage.

### SauUSI can change motor activity upon reaching a 5mC site

To further show that SauUSI interactions do not activate cleavage, we examined DNA with two hemimethylated sites. Depending on which strands are methylated, SauUSI motors will either travel in different directions (methylation on different strands - inverted repeats) or in the same direction (methylation on the same strand - direct repeats).

On linear substrates with excess SauUSI, cleavage rates were not significantly affected by relative site orientations, while at low concentrations, cleavage efficiency scaled as head-to-head (HtH) repeat >> tail-to-head (TtH) repeat > tail-to-tail (TtT) repeat (**Fig. 6A-C**). Since low concentration would not support multiple motors on each DNA, these results suggest that individual motors reaching a second site on HtH and TtH DNA leads to activation; on TtT DNA, motors travel towards DNA ends without reaching a second site. In the C-trap, DNA with sites in HtH repeat were more frequently cleaved than DNA with sites in direct repeat (**Fig. 4G**)

**Fig. 6.**
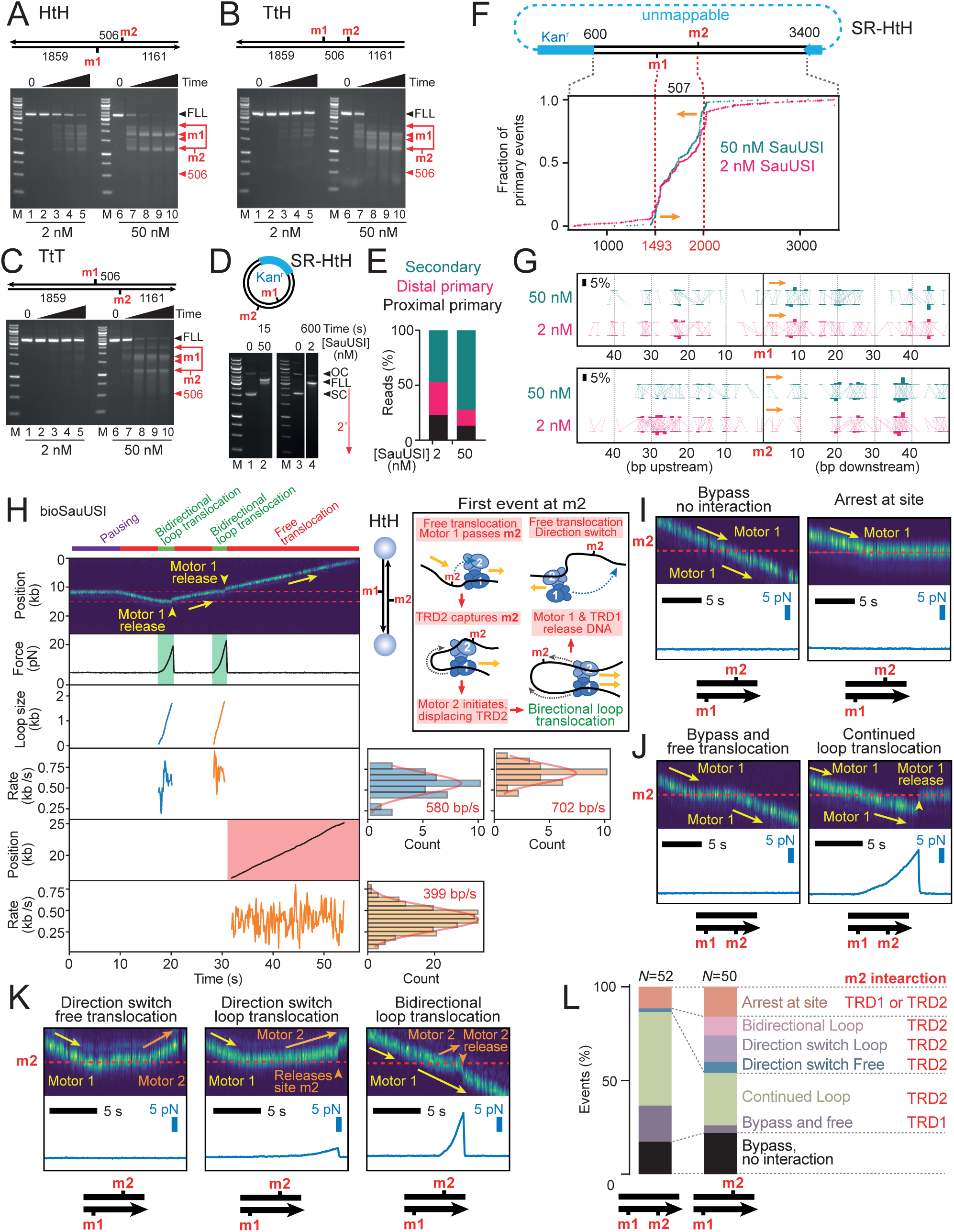
SauUSI motor activity can change upon reaching a 5-methyl cytosine site. **A-C**, Time courses of cleavage with sub-stoichiometric and excess SauUSI of 2-site DNA with sites in head-head (A), tail to head (TtH) and tail-to-tail (TtT) arrangements, analyzed by gel electrophoresis. Reaction times were 0, 0.25-, 5-, 10- and 20-min. **D**, Cleavage of HtH substrate ring by sub-stoichiometric SauUSI and excess SauUSI, analyzed by agarose gel electrophoresis. Cartoon shows SR-HTH map. **E**, Event types from panel D quantified using ENDO-Pore analysis. **F**, Cumulative percentage of primary events relative to the 5mC positions (m1 and m2) on SR-HTH. Yellow arrows indicate polarity of unidirectional translocation from each site. **G**, SLPs for each site oriented in the same translocation polarity with bars representing the percentage of phosphodiester cleavage at each top and bottom location and horizontal/vertical lines that indicate the ends generated (blunt or overhangs). **H**, Representative kymograph at 1.5 pN of a HtH DNA-bead tether with labelled enzyme (655 nm quantum dot-streptavidin-biotin-WT SauUSI) prebound at one 5mC site. Following a delay after moving into the ATP channel, free translocation is observed. Upon reaching the second 5mC (m2) that is on the non-translocated strand, an increase in force indicated bidirectional translocation (Cartoon). Collapse of the loop by motor 1 dissociation leads to free translocation back towards m1. Upon reaching m1, a force increase indicates bidirectional translocation that becomes free translocation upon motor 1 release. Quantum dot blinking produces occasional trace gaps. Instantaneous rates for loop or free translocation events were binned to give average rates from Gaussian fits (bin widths were 70, 65, and 50 bp/s for derivation of rates 580, 702, and 399 bp/s, respectively). **I-K**, example kymographs and force measurements that classify events – see main text for explanations. L, quantitation of event types from panels I-K where the second site is on the translocated or non-translocated strand. Red text indicates the suggested TRD interaction at m2 that produces the event.

Using ENDO-Pore to map HtH DNA with excess SauUSI, primary cleavage mainly occurred within the HtH arc consistent with convergent translocation (**Fig. 6F**). However, the absence of a strong peak near the midpoint (~1746 bp) suggests motor collisions do not trigger cleavage as seen for other translocating restriction enzymes^37,39,41,42^. Proximal events were mainly downstream of each 5mC (**Fig. 6G**), likely due to loop translocation across the short arc reaching the second site that triggers cleavage at the loop base (**Fig. 4H**).

With low SauSUI concentrations, cleavage still mainly occurred within the short HtH arc despite the reduced chance of collisions, suggesting that SauUSI can indeed recognize 5mCs on the non-translocated strand (**Fig. 6F**). Additional upstream proximal events (**Fig. 6G**) suggest bypass of the second site is also possible - there are also more distal events - and that translocation around the ring re-engages the original site, as seen with 1-site rings (**Fig. 4F**).

Using the C-trap, we directly observed how SauUSI motors interact with sites during free translocation on multi-site DNA. In the HtH example (**Fig. 6H**), SauUSI initiates free translocation at the first 5mC (m1). Upon reaching the second 5mC on the opposite strand (m2), increased force indicates bidirectional translocation. A force decrease due to Motor 1 releasing the loop leaves Motor 2 to free translocate back towards m1. Upon reaching m1, another bidirectional event was observed. A further force decrease marks Motor 1 dissociation, with Motor 2 resuming free translocation, in the opposite orientation to the original event.

While some translocation events bypassed the second site without measurable interaction, most showed evidence of 5mC recognition (**Fig. 6I–L**). After an interaction delay, “Bypass and free translocation” was more common when m2 was on the translocated strand (**Fig. 6J,L**), likely reflecting TRD1 binding and release without cleavage. If TRD2 binds m2 on the translocated strand, “continued loop translocation” occurs (**Fig. 6J**), as seen during hemimethylated loop initiation (**Fig. 2**). When m2 was on the non-translocated strand, TRD2 binding more often led to second motor loading, that then resulted in direction changes or bidirectional movement (**Fig. 6K**). “Arrest at site” reflects permanent stalling without cleavage activation for the duration of the experiment.

In summary, we do not find evidence that long-range interactions between SauUSI dimers activate cleavage. However, where motors encounter 5mCs, the ability to re-initiate translocation and in some cases to reverse direction allows a single enzyme to patrol back-and-forth between sites, increasing the chance of cleavage.

### Secondary cleavage results from cycles of site-binding, translocation and cleavage

After primary cleavage, SauUSI mediates secondary DNA shredding that increases with time as observed using gels and ENDO-Pore (**Fig. 4A–D**, **Fig. 5A**; **Extended data Fig. 4A**; **Fig. 6A-E**, **Extended data Fig. 5**). In ENDO-Pore, DNA shortening due to multiple breaks will be observed as pseudo 3ʹ-extensions with a spacing dictated by the outermost breaks^39^; SauUSI secondary cleavages were thus defined as 3ʹ extensions of >5 nt (**Fig. 4D**).

A wide range of 3ʹ spacings was observed even at early timepoints (**Fig. 4D**, **Extended data Fig. 5B**), indicating rapid accumulation of multiple random breaks—unlike the progressive end processing seen with Type I-SP restriction enzymes^39,43^—and with asymmetric distributions that reflect translocation directionality. Primary and secondary cleavages shared preferred loci, especially near the site (**Extended data Fig. 5C,D**), though later timepoints showed activity at a wider range of dinucleotides (**Supp Fig. 8B**). A lack of triplex displacement or cleavage of bystander non-methylated DNA suggests the motor and nuclease are inactivated after releasing the methylated DNA, so persistence of activated nucleases cannot explain shredding (**Supp Fig. 11**).

These results support a model of repeated 5mC binding, translocation, cleavage, and release (**Extended data Fig. 5D**), consistent with reduced shredding at low enzyme levels or when site-rebinding is blocked (**Fig. 4D**). Similar secondary distributions at low SauUSI or with M-Oligo (**Extended data Fig. 6**), suggest that loci are not favored by looped versus non-looped species. Our primary and secondary cleavage data do not support the previously proposed pile-up model but instead demonstrate that individual SauUSI dimers working alone can generate multiple dsDNA breaks.

## CONCLUSIONS

We describe a novel motor-driven cleavage mechanism in which binding to a modified site and translocation by a single SauUSI dimer triggers random dsDNA breaks. Alterations in motor activity upon rebinding 5mC is distinctive and enables continuous bidirectional scanning. While a single hemimethylated site is sufficient, clustered modifications on both stands that are more likely on plasmids and phages would promote repeated translocation and extensive cleavage, even at low enzyme levels. This shredding activity likely underlies the effectiveness of SauUSI in limiting HGT in *S. aureus*^18,19^. Crucially, cleavage is constrained *in cis*, ensuring specificity for methylated non-self DNA— unlike abortive systems such as Lamassu that release a diffusible nuclease to produce cell death^44–46^, SauUSI acts as a first-line defense, neutralizing threats without harming the cell and exemplifying the sophisticated strategies that have evolved for distinguishing invader from host.

For the Type I, I-SP, and III RM enzymes, ATP-driven long-range interactions are required to dimerize and activate nuclease subunits^5,28,29,42,43^. Such communication between ATP-dependent enzymes may have evolved to limit autoimmunity when self-DNA modifications are lost, for example, following DNA repair^47,48^. In contrast, SauUSI dimers are autonomous and likely cleave both DNA strands sequentially using the single PLD active site, akin to BfiI (Ref^27^). SauUSI motor activity could provide an additional HGT barrier by nucleoprotein remodeling, although we found that roadblock displacement was context dependent. However, a more likely role for movement on DNA is to promote random shredding of non-self DNA that will be hard to repair. SauUSI may anyway have lower metabolic cost as accidental host DNA cleavage would require the presumably less frequent horizontal acquisition of a cytosine methyltransferase.

Helicase domains are common in defense systems ^e.g. 49–52^, and their roles can extend beyond orthodox duplex unwinding, as shown here and elsewhere^38,53^. Upstream helicase-like motor initiation that displaces a TRD was also observed for the Type I-SP restriction-modification enzymes, but the translocating species did not form a loop^43,54,55^. Future work on SauUSI should explore the structural basis of loop translocation and of strand-swapping by its PLD nuclease domains during cleavage. The SauUSI motor activity may also shed light on the Type II restriction enzymes CglI and NgoAVII that also combine PLD nuclease dimers with dual helicase-like ATPases but which recognize specific unmodified sequence (e.g., 5’-GCSGC-3’ for CglI) ^56–58^. These systems share features with SauUSI—DNA translocation, cleavage near recognition sites, and activity on single-site DNA—but unlike SauUSI, can translocate onto adjacent DNA once activated. The coupling of motors and nucleases may prove to be a highly effective pairing in defenses by allowing transfer of nuclease activities away from target sites to introduce multiple random distantly-spaced toxic DNA breaks.

## METHODS

Unless stated otherwise, protocols followed the manufacturers’ recommended procedures. See Supplementary Excel file for oligonucleotide and gene sequences.

### Proteins

All protein purification methods are detailed in the Supplementary Methods.

### Methylation of DNA substrates

Methylation by M.AluI used equimolar methyltransferase to DNA with 640 µM *S*-adenosyl methionine (AdoMet) in AluI methyltransferase buffer (New England Biolabs, NEB). Methylation by M1 or M2.BfuAI used 20-fold higher concentrations than BfuAI sites with 800 µM AdoMet in M.BfuAI buffer (20 mM Tris (pH 7.4), 1 mM EDTA and 1 mM DTT). Reactions were incubated at 37 °C for 1 – 1.5 h before purification using AMPure XP beads (Beckman Coulter). Methylation levels were verified using AluI or BfuAI restriction endonucleases, as appropriate. Methylation by M.AluI was sequence-specific but both M1.BfuAI and M2.BfuAI were promiscuous, additionally methylating sites that differed by 1 bp from the cognate site (“star sites”), as seen for other bacterial MTases^59–62^. To avoid this, all DNA substrates methylated by M1 or M2.BfuAI had the star sites removed.

### DNA substrates

#### Substrate rings

To generate circular DNA, we used recombination by Tn*21* resolvase at pairs of *res* sites on recombinant plasmids to generate origin-free supercoiled DNA circles (Substrate Rings - SRs). Synthetic constructs pUC_1TB (Integrated DNA Technologies, IDT) and pUC_HtH (Twist Biosciences) were designed based on pSH1 (Ref^63^). Other constructs (pUC_1T, pUC_1B, pUC_HtT and pUC_TtT) were generated by site directed mutagenesis from these DNA using QuikChange or QuikChange Multi mutagenesis kits (Agilent). Methylation-free plasmids were purified from *E. coli* GM2929 (ref^64^) using a Qiagen Plasmid Maxi Kit. To generate a single-interlinked (−2) catenane, 20nM plasmid and 500 nM Tn*21* Resolvase were incubated in 50 mM Tris-Cl (pH 8.0), 20 mM potassium glutamate, 10 mM MgCl_2_ and 1 mM DTT for 1 h at 37 °C. Following purification (Zymo Clean & Concentrate kit), the non-SR DNA circle was linearized and fragmented with EcoRI-HF (NEB) and 10 nM RecBCD (gift from Mark Dillingham) in rCutSmart buffer (NEB) supplemented with 1 mM ATP for 1.5 h at 37 °C. SRs were purified using AMPure XP beads (Beckman Coulter). SR1TB was methylated by M.AluI whereas all other SRs were methylated by M1.BfuAI, as above.

#### Linear DNA for biochemical assays

For cleavage assays, SR1B was digested with SpeI, NcoI or PshAI (NEB) to generate S1B, N1B and P1B, respectively, while linear HtH, HtT and TtT substrates were generated by cleavage of the relevant SR with NcoI (NEB). To generate linear DNA with hairpin ends, SR1B was amplified by PCR to incorporate BsaI sites at both ends. Two DNA hairpins with BsaI sites complementary to each end were generated from oligonucleotides (IDT) by heating to 95 °C for 5 min in 10 mM Tris, 1 mM EDTA and 50 mM NaCl before gently cooling to room temperature. 5 nM linear DNA and 125 nM of each loop were incubated in T4 DNA Ligase buffer (NEB) with 1/20 volume of BsaI/ligase mix (NEB) followed by thermal cycling for >100 cycles at 10 °C for 30 s followed by 37 °C for 30 s, with ramping of 0.2 °C/s between cycles. The sample was supplemented with 1 mM ATP and treated with 10 nM RecBCD (gift from Mark Dillingham) for 1 hour at 37 °C prior to heat inactivation at 65 °C for 10 min.

For the PBP assays, linear DNA was generated by PCR using SR1B and the methylated version produced using M1.BfuAI, as above.

Dual TFO M.AluI substrates and the 53, 103, 153 and 203 bp spacings were generated by PCR from synthetic DNA (IDT). The Dual TFO BfuAI variant was generated from the AluI variant by QuikChange mutagenesis (Agilent). Triplex substrates with variable upstream DNA were produced by annealing synthetic DNA (IDT). The dCas9 triplex substrate was generated by introducing a triplex binding site into using amplification of SR1B with primers that introduced the TFR. DNAs were methylated as above.

#### C-trap

Biotinylated handles were produced by incubating Lambda DNA (*dam-*, *dcm-*, ThermoFisher Scientific) with Klenow fragment (3ʹ→5ʹ exo-, NEB), 50 μM dATP/dTTP/dGTP, and 40 μM biotin-14-dCTP in NEB Buffer 2 at 37 C for 30 min. After heat inactivation at 75 C for 15 min, the DNA was digested with BsaI-HFv2 (NEB) at 37 C for 3 h and the 11.4 kb fragment purified following agarose gel electrophoresis using QIAEX II beads (Qiagen). A 34 bp hemimethylated insert with appropriate 5’ overhangs was annealed from synthetic DNA (IDT). All other inserts were amplified from SRs using primers containing BsaI sites to produce 3.4 kb products that were BsaI-digested and methylated as above. Handles and inserts were ligated at a 2:1 molar ratio with T4 ligase overnight at 16 °C and purified using AMPure XP beads (Beckman Coulter).

#### Magnetic tweezers

Using either the SR1B or SR1TB, linear DNAs with BsaI restriction sites at each end were generated by PCR and methylated as above. 1.0 kb biotin-or digoxigenin-modified attachment handles were made by PCR using pUC19 (ref^65^) with biotin- or digoxigenin-dUTP (Sigma Aldrich) using primers including BsaI sites that were complementary to one or other end of the substrate DNA. Handles and substrate DNA were assembled with a single site-specific nick using a one-pot Golden Gate protocol: (1) 112.5 nM biotin-handle in 20 µL was cleaved and dephosphorylated with FastDigest Eco31I (a BsaI isoschizomer) and Fast Alkaline Phosphatase (both ThermoFisher Scientific) for 10 min at 37 °C followed by 15 min at 65 °C; (2) 25 nM DIG-handle, 2 nM insert, and 4.5 uL BsaI/ligase mix (NEB) and 9 µL NEB ligation buffer (NEB) were added, in a final reaction of 90 µL; (3) The reaction was incubated in a thermal cycler for >100 cycles of 10 °C for 30 s and 37 °C for 30 s with 0.2 °C/s ramping between cycles; (4) Following incubation at 67 °C for 10 min, the DNA with both handles attached was purified following agarose gel electrophoresis in the absence of DNA dyes using QIAEX II beads (Qiagen).

#### Oligoduplex traps

To make 69 bp oligoduplex traps, synthetic DNA (IDT) were annealed: Sau_R (5ʹ-GCATAGAAATTGCATCAACGCATATAGCGCTAGCAGCACCGCCATAGTGACTGGCGATGCTGTCGGAAT-3ʹ) was annealed with Sau_FM (5ʹ-ATTCCGACAGCATCGCCAGTCACTATGGCGGTGCmTGCTAGCGCTATATGC GTTGATGCAATTTCTATGC-3ʹ, where Cm is 5-methylcytosince) to make a hemimethylated trap (M-Oligo); or Sau_R was annealed with Sau_F (5ʹ-ATTCCGACAGCATCGCCAGTCACTATGGCGGTGCTGCTAGCGCTATATGCGTTGATGCAATTTCTATGC-3ʹ) to make a non-methylated control trap (NM-Oligo).

### Triplex displacement assays

#### Labelling

For gel-based assays, Triplex Forming Oligonucleotides (TFOs) were 5ʹ-labelled with either Cyanine 3 (IDT) or ^32^P. For the latter, TFOs were labelled using T4 polynucleotide kinase and [γ-^32^P]ATP at 37 °C for 1 hr in T4 PnK Buffer (New England Biolabs). This was followed by a further incubation at 80 °C for 10 min before purification using Micro Bio-Spin chromatography columns (Bio-Rad Laboratories). In stopped flow assays, TFOs were 5ʹ-labelled with 5-Carboxytetramethylrhodamine (IDT). Triplexes were formed by heating 2:1 duplex DNA to TFO in 50 mM MES (pH 5.5) and 15 mM MgCl_2_ at 57 °C for 15 min before slowly cooling to ambient temperature.

#### Gel-based assays

10 nM Triplex DNA was incubated with 25 nM SauUSI dimer in Reaction Buffer (RB) (50 mM Tris-Cl (pH 8.0), 100 mM NaCl, 10 mM MgCl_2_, 1 mM DTT and 100 µg ml^-1^ bovine serum albumin (BSA)) and reactions initiated by addition of 4 mM ATP (where indicated). Reactions were incubated at 25 °C for 20 min before the addition of 5x FMS (12.5 % (w/v) Ficoll-400, 250 mM MOPS (pH 5.5) and 3% (w/v) SDS). For “melted triplex” controls, the samples were heated at 80 °C for 5 min followed by the addition of 5x FMS. Samples were separated on 5% (w/v) polyacrylamide (19:1) gels containing TAM (40 mM Tris-Acetate (pH 7.0) and 1 mM MgCl_2_) at 12.5 V/cm at 4 °C. For ^32^P-labelled triplexes, gels were dried onto 3MM Chr blotting paper (Whatman) at 80 °C for 1 hr under vacuum using a Model 583 Gel Dryer (Bio-Rad Laboratories). Dried gels were visualized using a Typhoon FLA 9500 (Cytiva). For Cy3-labelled triplexes, gels were visualized without drying using the Typhoon FLA 9500. Band intensity was quantified using ImageJ v. 2.16.0 (National Institutes of Health).

#### Stopped flow assays

Stopped flow experiments used a TgK Scientific SF-61 DX2 double mixing stopped-flow system in single mixing mode with temperature control (±0.1 °C) via a water bath-fed manifold around the syringes and observation cell. A 570 nm long-pass filter was used with excitation wavelength (75-W mercury-xenon lamp) set to 548 nm. Reactions were mixed as indicated with final concentrations of 1 nM triplex DNA, varying concentrations of WT SauUSI dimer, and 0 - 4 mM ATP in RB. ATP dependence experiments additionally had 25 mM phosphocreatine (Sigma Aldrich) and 5 U/mL creatine phosphokinase (Sigma Aldrich) to maintain ATP concentrations. Where included, traps were at final concentrations of 5 µM Heparin, 1 µM M-Oligo or 0.25 µM K255A helicase mutant. Temperature dependence assays were performed at 20 - 37 °C by mixing 1 nM triplex DNA and 25 nM WT SauUSI dimer in RB with 4 mM ATP.

#### Stopped flow analysis

Time traces were recorded for >1 minute, until complete triplex displacement was observed. Data from 3 traces were averaged using Kinetic Studio 5.1.0 (TgK Scientific). Each experiment was repeated at least twice and data analyzed using Graphpad Prism Ver 10.1. Lag times were determined by fitting to the incorporated “Plateau followed by single exponential” equation. Lag times were used to calculate the observed translocation rate by linear regression. Triplex displacement burst sizes in the presence of trap were determined by fitting to the “Plateau followed by double exponential” equation.

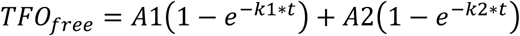

Where TFO_free_ is the displaced triplex, *A*1 and *k*1 are the amplitude and rate, respectively, of the first phase and *A*2 and *k*2 are the amplitude and rate, respectively, of the second phase.

For temperature dependence experiments, lag times were measured in duplicate with four spacings and fitted to give rates that were analyzed using:

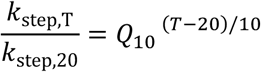

Where *k*_step,T_ is the stepping rate at temperature T, *k*_step,20_ is the rate at 20 C, and *Q*_10_ is a temperature coefficient that gives the fold-increase in rate for a 10 C increase assuming no denaturation effects.

### ATPase assays

#### Measuring phosphate release using Phosphate Binding Protein

Experiments were performed in the SF-61 DX2 with single mixing at 25 °C. A 455GG filter was used to record MDCC emissions with the excitation wavelength set to 437 nm. 7-Diethylamino-3-((((2-maleimidyl)ethyl)amino)carbonyl)coumarin-labelled Phosphate Binding Protein was calibrated by titrating inorganic phosphate (Pi) against 12 µM PBP and recording the emissions for 15 s with 2048 points (**Supp. Fig. 12**). SauUSI reactions were mixed as indicated with final concentrations of 0.5 nM 1B substrate (methylated or unmethylated), 12.5 nM WT SauUSI dimer, 0 - 4 mM ATP, 12 µM PBP and 0.25 µM SauUSI K255A dimer, 0.05 U mL^-1^ bacterial purine nucleoside phosphorylase (Sigma Aldrich) and 0.2 mM 7-methylguanine (Sigma Aldrich) in RB.

#### ATPase analysis

PBP calibration was determined by fitting the average photomultiplier signal at each Pi concentration by linear regression in GraphPad Prism V10.1 (**Supp. Fig. 12**). Pi release due to ATP hydrolysis by non-specifically bound SauUSI was removed by subtracting data measured with unmethylated DNA from data measured with methylated DNA. Traces were converted to “ATP hydrolyzed per DNA” by dividing by the DNA (5mC) concentration. Initial ATPase rates were determined by linear regression. As only a percentage of SauUSI WT successfully initiates translocation, the ATPase rates were adjusted using the first burst phase amplitude (*A*1) from the triplex displacement data, above. The adjusted *k*_ATP_ values were fitted to a Michaelis-Menten equation in GraphPad Prism to determine *V*_max_ and *K*_M_.

### Topoisomerase assay

Relaxed SRs were produced by incubating 5 μg DNA with 8U Wheat Germ Topoisomerase I (Inspiralis) in NEB buffer r2.1 at 37 °C for 30 min, followed by purification with AMPure XP beads (Beckman Coulter). Topoisomerase supercoiling assay reactions contained 6.9 nM relaxed DNA, 75 nM SauUSI H119A, and 0.2 U/μL *E. coli* topoisomerase I (NEB) in NEB CutSmart buffer. An initiation mix of ATP alone or combined with trap (M-Oligo or SauUSI K255A) was made up in CutSmart buffer and reactions started by adding this mix to a final ATP and trap concentration of 4 mM and 1.5 μM, respectively. Reactions were incubated at 37 °C for 30 min and separated by agarose gel electrophoresis without ethidium bromide. Gels were imaged using a EpiChem^3^ Darkroom (UVP bioimaging) using 326 nm illumination.

### DNA cleavage assays

5 nM DNA with SauUSI concentrations indicated was preincubated in RB at 25 °C and reactions initiated by adding ATP to 4 mM (plus 1 µM M-Oligo or SauUSI K255A, where indicated). Reactions were quenched with 0.2 volumes DNA:Protein loading dye (10 mM Tris (pH 8.0), 100 mM EDTA, 15% (v/v) Ficoll-400, 1% (w/v) SDS, 0.03% (w/v) Bromophenol blue and 0.03% (w/v) Xylene Cyanol FF). Samples were separated on by agarose gel electrophoresis and imaged as above.

### Cas9 roadblocks

gRNAs were produced using the EnGen sgRNA Synthesis Kit (NEB) and mixed 2:1 with WT Cas9 or dCas9 (NEB) in RNase-free SB (10 mM Tris-Cl (pH 7.5), 100 mM NaCl, 1 mM EDTA, 0.1 mM DTT and 5 µg/mL BSA) before incubation at 37 °C for 1 h. For dCas9 triplex displacement experiments, 10 nM Cy3-triplex DNA was preincubated with 100 nM dCas9 at 25 °C for 20 min. 25 nM SauUSI dimer was added where appropriate and reactions initiated by 4 mM ATP or 50 nM WT Cas9, and incubated for 20 min at 25 °C before addition of 0.2 vol of 5 x FSB. Samples were separated using 5% (w/v) acrylamide (19:1) in TAM for 1 hr at 12.5 V/cm. Gels were imaged with a Typhoon FLA 9500 (Cytiva).

For cleavage reactions with dCas9 roadblocks, 5 nM DNA was pre-incubated with 100 nM dCas9:gRNA for 20 min at 25 °C. 25 nM SauUSI was added if required and reactions initiated by either 4 mM ATP (and 1 µM M-Oligo, where indicated) or 50 nM WT Cas9. Samples were incubated at 25 °C for 10 min before quenching with 0.2 vol 5x DNA:Protein loading dye. Samples were separated by agarose gel electrophoresis and imaged as above.

### ENDO-Pore mapping of double-strand DNA breaks

ENDO-Pore was performed as previously described^39^. In brief, SauUSI reactions were quenched in Binding Buffer from the Clean and Concentrate kit (Zymo Research) and purified. Samples were treated with Ultra II end repair mix (NEB). A pUC19 origin sequence^65^ was amplified by PCR and treated with XcmI (NEB) to generate dT overhangs and this was ligated to the end repair samples using T4 DNA Ligase (NEB). *E. coli* OmniMAX™ 2-T1R cells (ThermoFisher) were transformed with each sample and single colonies selected after overnight incubation on LB agar with 50 µg/mL kanamycin. >40,000 colonies were scraped and directly purified using HiSpeed Plasmid Midi Kit (Qiagen). The DNA was subjected to rolling circle amplification using EquiPhi29 DNA polymerase with random primers (Thermo Fisher Scientific) to produce long concatemeric repeats before debranching by T7 Endonuclease I (NEB). Following size exclusion using the SRE XS kit (PacBio), the DNA was sequenced using Oxford Nanopore Minion R10.4.1 flow cells.

#### Analysis

Raw squiggle data was basecalled and demultiplexed using Dorado (https://github.com/nanoporetech/dorado/ V1.1.1). Concatemeric consensus was called for the demultiplexed output files using C3POa with the pUC19 origin as the specified repeat sequence (https://github.com/christopher-vollmers/C3POa V3.2). Consensus FASTA outputs were then analysed using Cleavage Site Investigator (CSI) with a >3 repeat filter (https://github.com/sjcross/CleavageSiteInvestigator v1.0.0). CSI output data was analyzed using Excel 365 (Microsoft) and GraphPad Prism v10.3.

### Combined optical tweezers and confocal microscopy (C-Trap)

Experiments used a LUMICKS C-Trap G2 system integrating a dual optical trap and confocal microscope and C1-type microfluidics flow cell. The imaging buffer used in all flow cell channels (Ch1-5) was 50 mM Tris-Cl (pH 7.9), 50 mM NaCl, 10 mM MgCl2, 0.01% (v/v) Tween-20 and 0.2 mg/ml BSA. Biotinylated WT or H119A SauUSI (1 nM) was incubated with 2 nM streptavidin-conjugated 655-nm quantum dots (Qdots, Invitrogen) for 3 min at room temperature. Biotin was added to 1 μM to block free streptavidin Qdot sites before flowing this mixture into channel 4. Streptavidin-coated polystyrene beads (4.0–4.9 μm diameter, LUMICKS) were trapped by two 1,064 nm lasers (25% power) and used to form single DNA tethers under flow. The applied DNA force was set to 1.5 pN in Ch3 before moving the tether into Ch4 to bind Qdot-labelled SauUSI. Finally, the DNA and captured enzyme(s) were moved into Ch5 containing 4 mM ATP to visualize motor-dependent enzyme dynamics. Qdots were excited with a 488 nm laser at 2% power and imaged with a 680/42 nm emission filter as kymographs along the tether axis using a pixel exposure time of 0.1 ms, line scan time of 28 ms and pixel size of 100 nm.

#### Analysis

Raw kymograph and force data were analyzed in Jupyter notebooks using Python 3. The primary packages used were LUMICKS Pylake v.1.7.0, ctraptools (https://github.com/sjcross/ctraptools v0.3.2), scipy, pandas, and matplotlib. Free translocation rates were determined by first tracking Qdot trajectories using the Pylake KymoWidgetGreedy widget followed by track refinement with refine_tracks_gaussian and smoothing with a moving average filter. The first-time derivatives of track positions were then binned and their mean and standard deviation determined from a Gaussian fitted to this histogram using scipy.optimize.curve_fit.

To extract loop translocation rates, force data for individual looping events was manually selected to include several seconds of invariant baseline force before initiation and after loop collapse. Force drift was then accounted for by subtracting a second order polynomial fitted to this baseline data (where F_baseline_<F_min_+0.05*(F_max_-F_min_)). The loop translocation event was then classified as contiguous time points for which F > 〈F_baseline_〉+2*σ_baseline_. Force was converted into loop size in base pairs as described below, allowing calculation of instantaneous translocation rate for each timepoint as the first time-derivative. After manually excluding data points corresponding to motor slippage, mean looping rates, for each event were assigned by binning instantaneous rates and fitting a Gaussian. Translocation rate distributions were compared in GraphPad Prism v10.3 using an unpaired t-test with Welch’s correction for unequal variances.

### Magnetic Tweezers (MS)

Single-molecule experiments were carried out as previously^43,67,68^, using a commercial magnetic tweezers microscope (Picotwist) and PicoJai (v2019) software, with images acquired at 60 Hz with a Jai CV-A10 GE camera. Substrate DNAs were tethered to 1 µm MyOne paramagnetic beads (Invitrogen) and anchored in the anti-digoxigenin-coated flow cells using established methods^68^. Non-magnetic beads attached to the coverslip surface^69^ were monitored to correct for instrument drift. The 3D magnetic bead position and thus the length of the attached DNA was determined from video images using real-time 3D particle tracking with sub-nm accuracy^67^. Raw apparent DNA length data collected at 60 Hz was filtered using a non-linear filter^70^ in PIAS (vMarch 2024, PicoTwist). To determine the instantaneous rates of each event, the first order derivative was calculated and a 0.1 Hz median filter applied using Origin 2024b (OriginLab Corporation).

### Statistical analysis and reproducibility

The *n* values for the number of events are stated in each figure where relevant. Each single-molecule experiment was carried out on at least three different DNA molecules. Gel-based assays were repeated at least twice, with representative examples shown.

## Reporting summary

Further information on research design is available in the Nature Portfolio Reporting Summary linked to this article.

## Data availability

Example data for the ensemble and single-molecule and ensemble assays are presented within the paper. The full datasets that support the findings of this study are available at the University of Bristol data repository (to be deposited before publication).

## Code availability

LUMICKS Pylake is available at https://github.com/lumicks/pylake

The ctraptools package is available at https://github.com/sjcross/ctraptools

AlphaFold 3 is available at https://alphafoldserver.com/

Dorado is available at https://github.com/nanoporetech/dorado/

C3POa is available at https://github.com/christopher-vollmers/C3POa

Cleavage Site Investigator is available at https://github.com/sjcross/CleavageSiteInvestigator

WolframAlpha is available at https://www.wolframalpha.com/

ImageJ is available at https://imagej.net/ij/

## Supporting information

Oligonucleotide and gene sequences

## Acknowledgements

We thank Sophia Taylor and Alan Scott for preliminary experiments and Mark Dillingham and Tom Chambers for supplying purified RecBCD. This work was supported by the European Research Council under the European Union’s Horizon 2020 research and innovation program (ERC-2017-ADG-788405 to M.D.S. and by the BBSRC (21ALERT BB/W019337/1 to M.D.S.; sLoLa BB/X003051/1 to M.D.S.). For the purpose of open access, the authors have applied a Creative Commons Attribution (CC BY) license to any author accepted manuscript version arising.

## Author contributions

M.D.S. conceptualized the study. S.J.S., A,H-G., O.E.T.M., F.M.D. purified and labeled recombinant proteins and DNA. A,H-G., S.M., S.J.C. and M.D.S. designed, performed and analyzed single-molecule experiments. S.J.S., A,H-G., O.E.T.M. and D.D. designed, performed and analyzed data from the ensemble assays. S.J.S., O.E.T.M. and M.D.S. designed, performed and analyzed data from the ENDO-Pore assays. All authors contributed to the original draft and reviewed and edited the paper.

---

**Extended Data Fig. 1.**
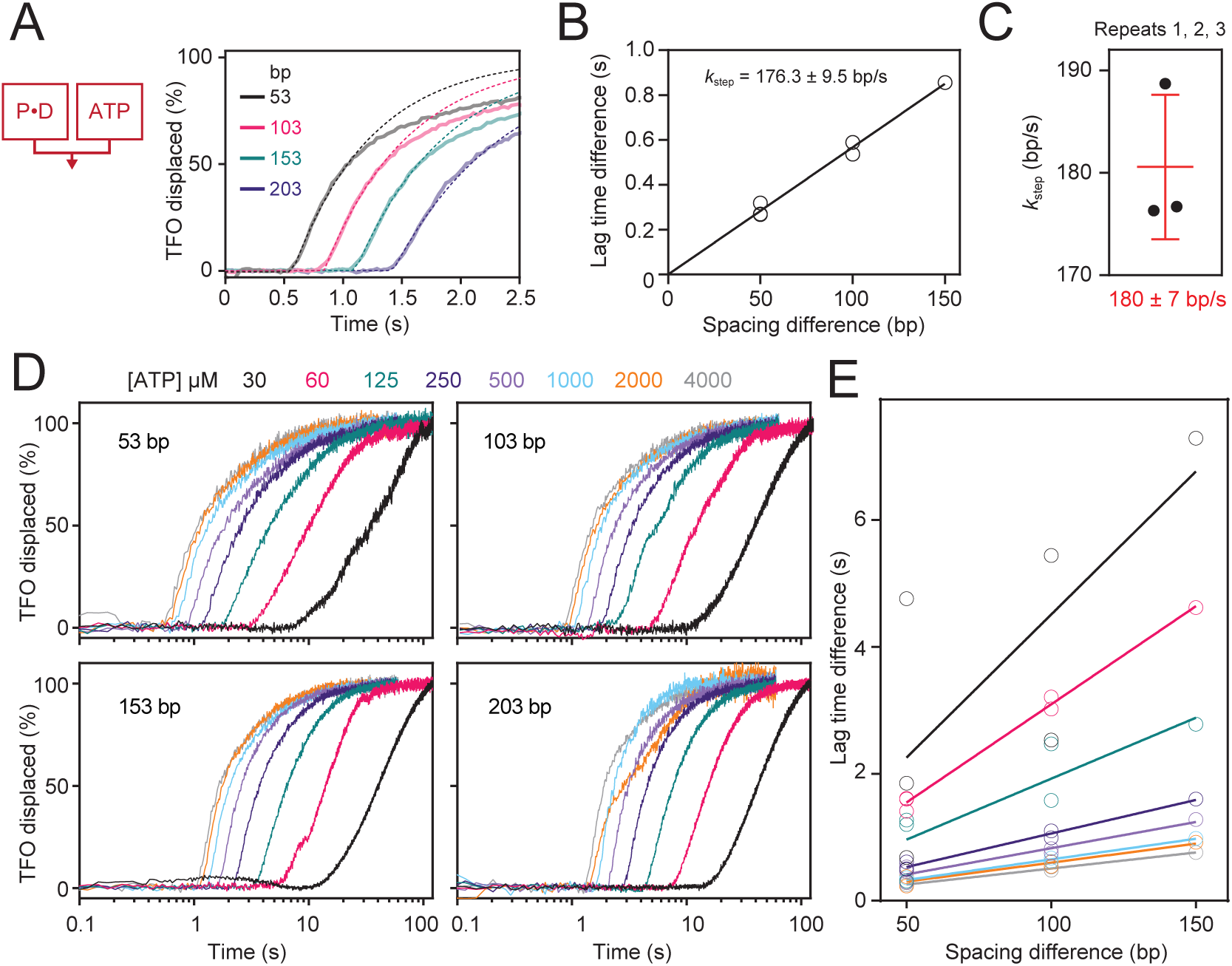
DNA translocation rates from triplex displacement assays. **A**, One repeat from a time-dependent triplex displacement from linear DNA with different hemimethylation – triplex distances (**Fig. 1C**) measured using a stopped flow assay. Samples were mixed as indicated (cartoon) with Protein (P), DNA (D) and ATP. Fits (dotted lines) were to a plateau followed by one phase association, with the plateau giving the lag time. **B**, Lag times from the repeat in Panel A were used to calculate the lag time differences for different spacing differences to give six values (some overlapping in the figure). Linear regression fixed to pass through the origin gives the translocation rate with standard error. **C**, Translocation rates from three repeats, with the mean and standard deviation. **D**, One repeat from a time-dependent triplex displacement from linear DNA with different hemimethylation – triplex distances and ATP concentrations, measure as in Panel A. **E**, Lag times calculated from Panel D were used to determine linear fits to the lag time differences against spacing differences as in panel B. Translocation rates from both repeats are plotted in **Fig. 1F**.

**Extended Data Fig. 2.**
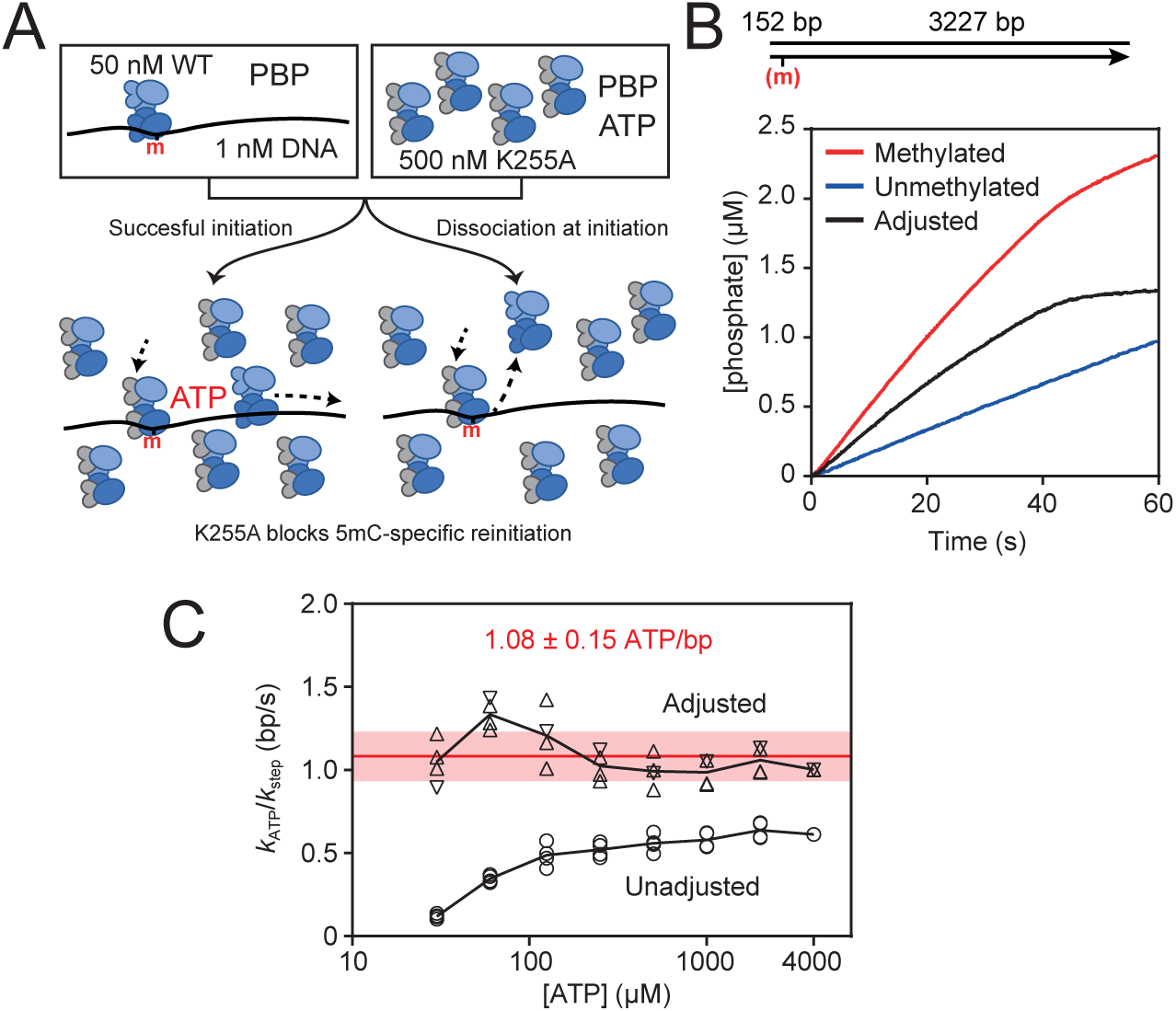
ATPase assays using phosphate binding protein. **A**, Model mixing scheme for the stopped flow ATPase assays with phosphate binding protein (PBP). The K255A helicase mutant was used as a trap to prevent site rebinding following initiation. **B**, Stopped flow phosphate release data using the methylated and unmethylated versions of the substrate shown. The background ATPase hydrolysis due to non-specific binding was subtracted to obtain the 5mC-specific rate. **C**, Four estimates for the coupling ratio (*k*_ATP_/*k*_step_) calculated from the two ATPase and two triplex repeats. For the adjusted values, the ATP-dependent initiation bursts from the triplex assay (**Fig. 1E**) have been used to correct *k*_ATP_ to account for the concentration of translocating species being lower than the 5mC concentration due to initiation failure. Mean (red line) and SD (red rectangle) coupling value is shown from the adjusted values.

**Extended Data Fig. 3.**
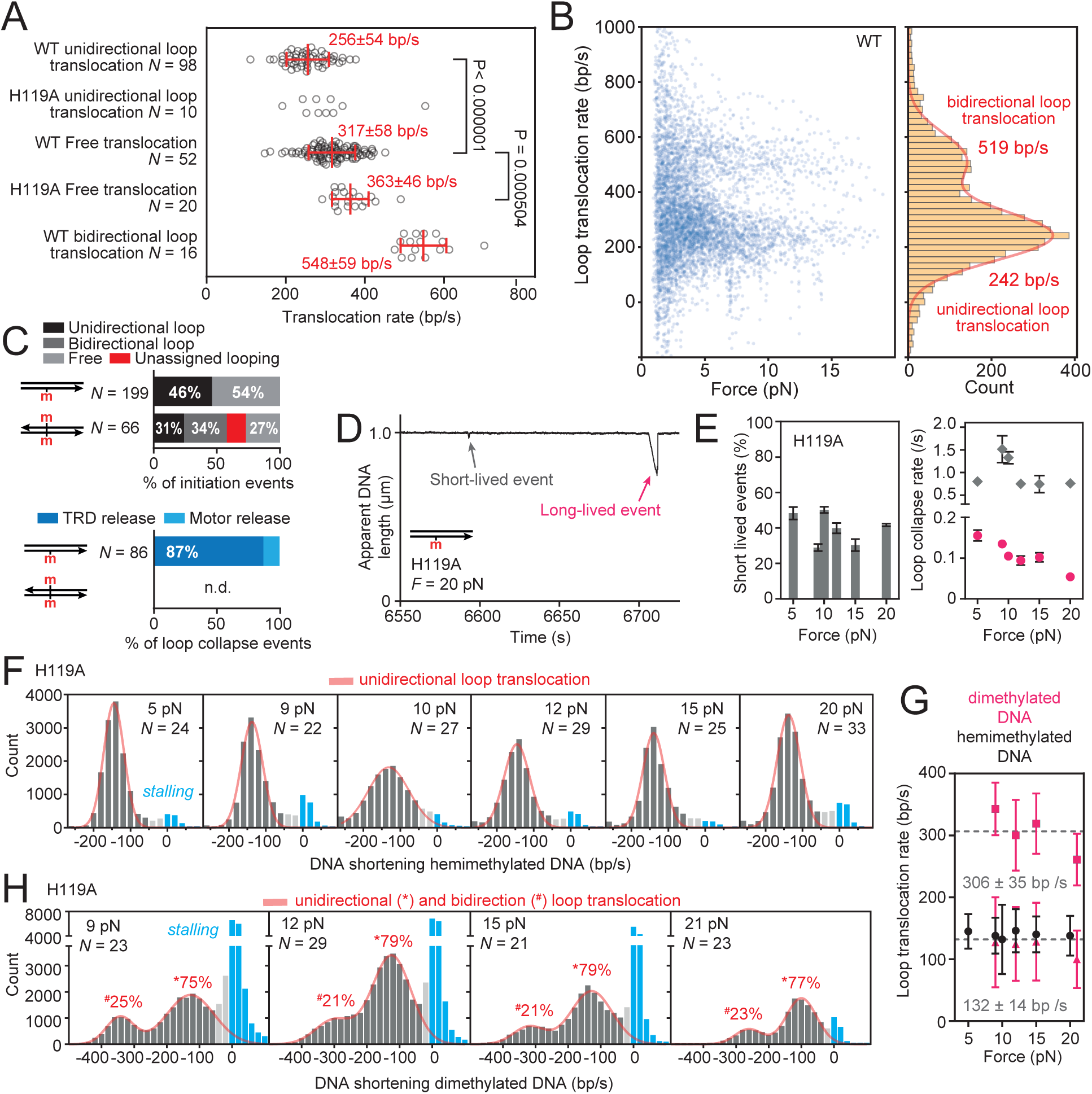
Single-molecule DNA translocation parameters. **A**, Translocation rates for individual events calculated from Gaussian fits to the instantaneous rates (e.g. **Fig. 2A**). Red lines and values show the mean and standard deviation. P values were from an unpaired t-test with Welch’s correction for unequal variances. **B**, Relationship of instantaneous rates to force for all observed events. Rates were binned (width = 22 bp/s) and fitted to a double Gaussian to give average rates for uni- and bidirectional rates. **C**, Percentages of loop versus free initiation events and TRD or motor release during loop collapse events. Events on dimethylated DNA were generally too complex to confidently assign the loop collapse event. **D**, Example MT trace of apparent DNA length showing short- and long-lived events for SauUSI H119A. **E**, Force-dependence of loop collapse rates for short-lived (<1s) and long-lived (>1s) events (*right panel*), and percentages of short-lived events (<1s duration, *left panel*). **F**, Instantaneous loop translocation rates (20 bp/s) at different DNA stretching forces using hemimethylated DNA. Blue events represent stalling during translocation. Gaussian fits (red) omitted data close to the stalling events (light grey). **G**, Force-dependence of the average loop translocation rates using hemi- or demethylated DNA. Grey lines are the average rates for uni- and bidirectional events. **H**, Instantaneous loop translocation rates (20 bp/s) at different DNA stretching forces using demethylated DNA. Fits to the sum of two Gaussians (red) omitted data close to the stalling events (light grey). Percentages were calculated from the area under each Gaussian estimated from the fits. Error bars in all panels are standard deviation.

**Extended Data Fig. 4.**
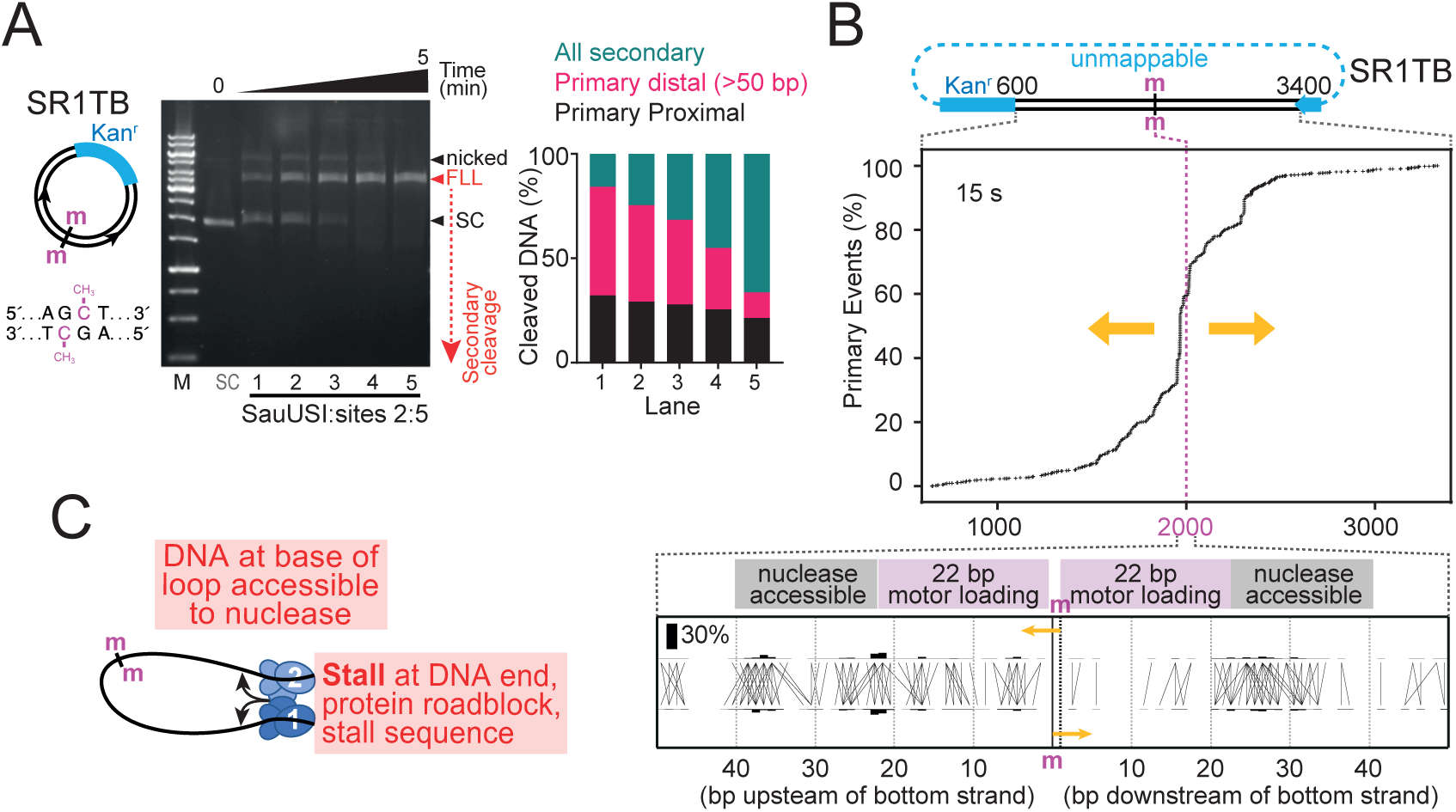
Dimethylated DNA cleavage is influenced by bidirectional translocation. **A**, Time-course of cleavage of dimethylated substrate ring SR1TB by sub-stoichiometric SauUSI, measured using agarose gel electrophoresis. Cartoon shows SR1TB map. Event types were quantified from ENDO-Pore analysis of the time points (0, 0.25, 0.5, 1, 2, 5 min). Primary cleavage is more efficient than on SR1B under equivalent conditions. **B**, Cumulative percentage of primary events relative to the 5mC positions (bottom strand 2000 and top strand 2001) on SR1TB. Yellow arrows indicate bidirectional translocation. For the proximal region, a strand linkage plot (SLP) shows bars representing the percentage of phosphodiester cleavage at each top and bottom location and horizontal/vertical lines that indicate the ends generated (blunt or overhangs). **C**, Model for DNA cleavage during bidirectional loop translocation to give only distal cleavage.

**Extended Data Fig. 5.**
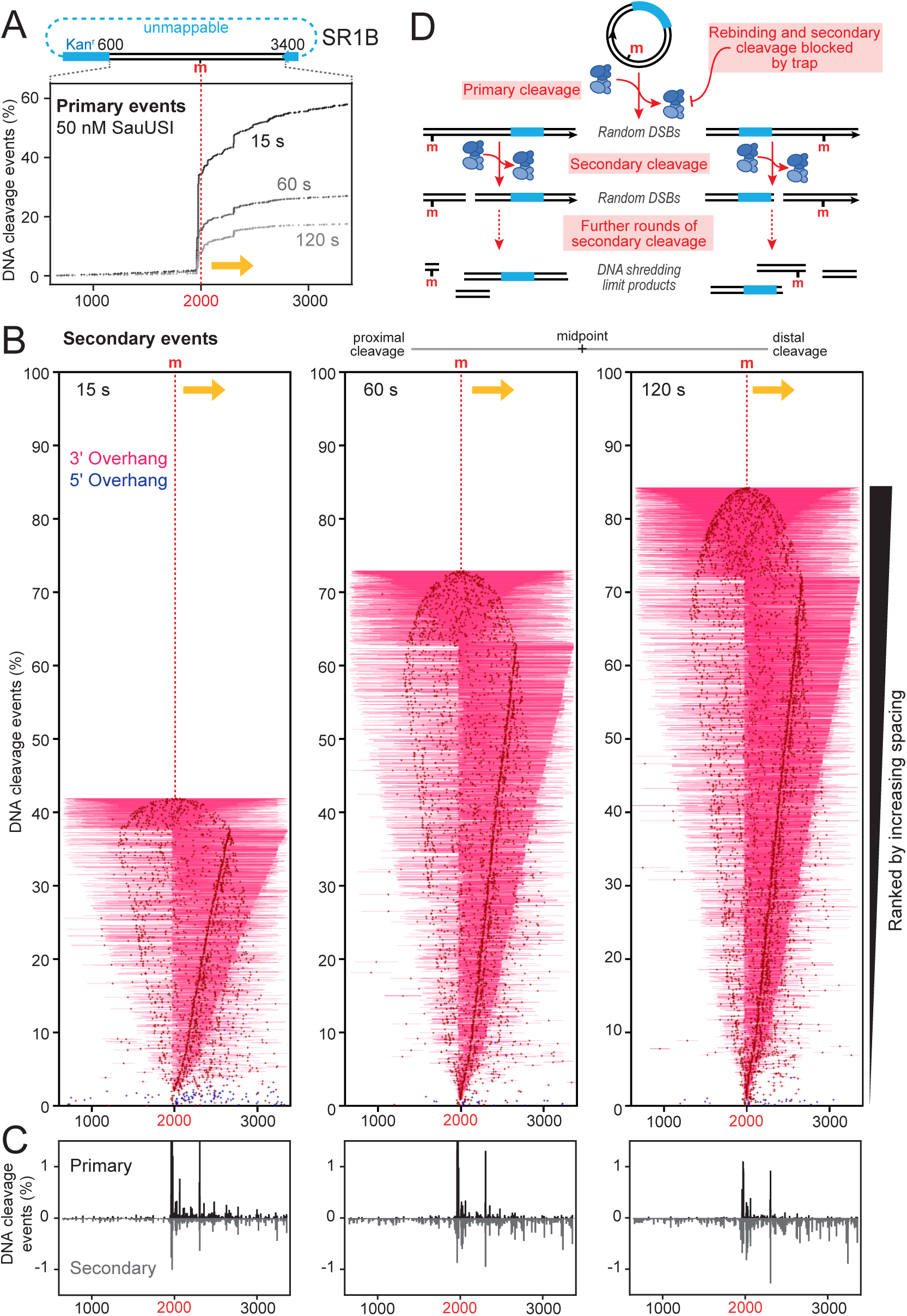
Secondary cleavage results from multiple translocation-cleavage events. **A**, Cumulative percentage of primary events relative to the 5mC position (m) on SR1B for the 15, 60 and 120 s reactions with excess (50 nM) SauUSI. Cleavage within the Kanamycin resistance cassette cannot be mapped by ENDO-Pore. Yellow arrow indicates polarity of unidirectional translocation. **B**, Secondary cleavage tornado plots where the crosses indicate the midpoint between outermost cleavages and the horizontal lines terminate at the outermost cleavage loci, with the events ranked by size and by position from left to right on the map. **C**, Positions of top and bottom strand primary cleavages (positive values) or secondary cleavages (negative values), binned with 1 nt bins. **D**, Model for production of secondary cleavage events. See text for full details.

**Extended Data Fig. 6.**
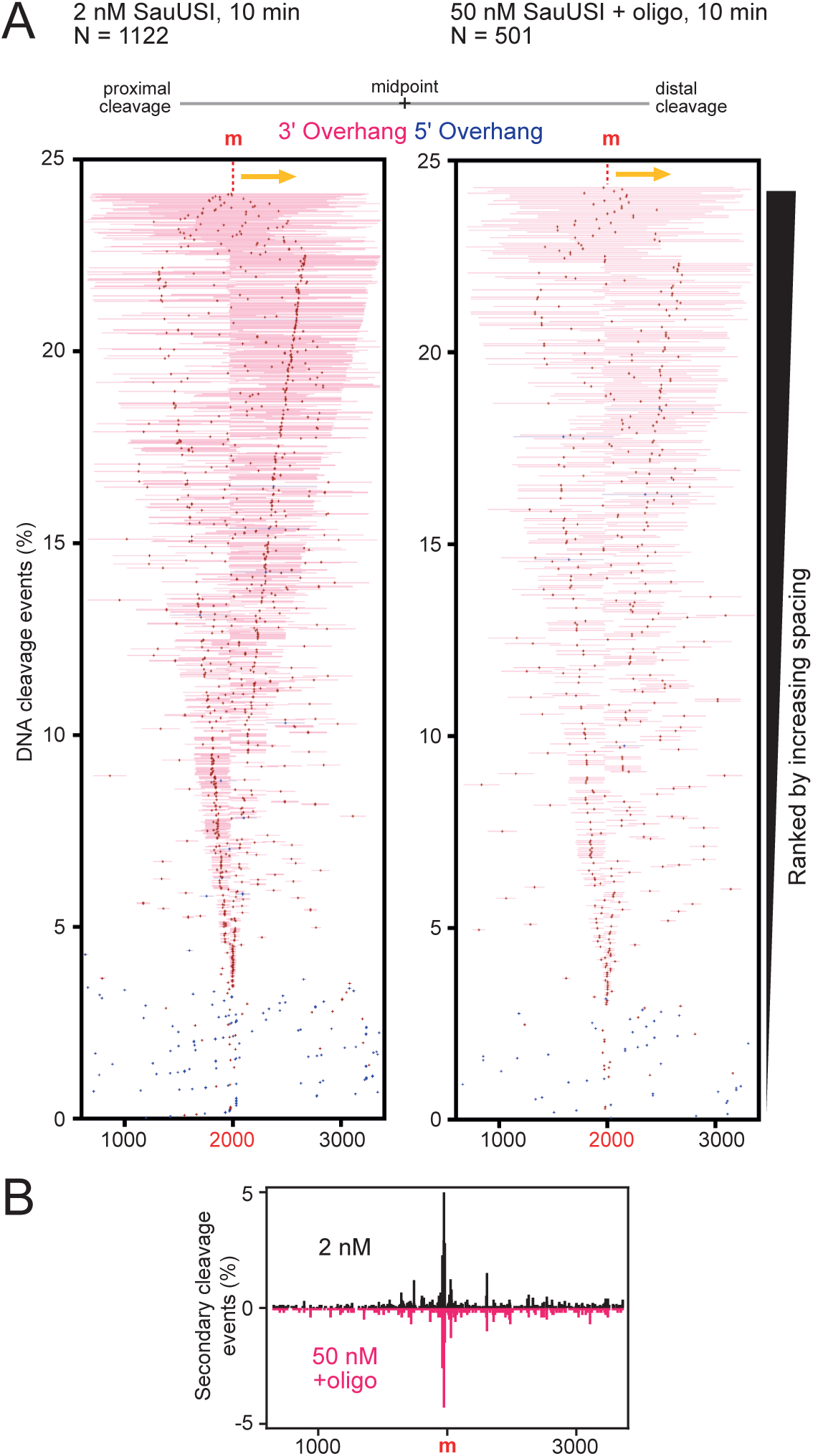
Cleavage loci are not determined by looping. **A**, Secondary cleavage tornado plots for low SauUSI or excess SauUSI with oligo trap, where the crosses indicate the midpoint between outermost cleavages and the horizontal lines terminate at the outermost cleavage loci, with events ranked by size and by position from left to right on the map. **C**, Positions of top and bottom strand secondary cleavages for low SauUSI (positive values) or excess SauUSI with oligo trap secondary cleavages (negative values), binned with 1 nt bins.

## REFERENCES

1. Isaev, A.B., Musharova, O.S. & Severinov, K.V. Microbial Arsenal of Antiviral Defenses - Part I. Biochemistry-Moscow 86, 319–337 (2021).

2. Isaev, A.B., Musharova, O.S. & Severinov, K. Microbial Arsenal of Antiviral Defenses. Part II. Biochemistry-Moscow 86, 449–470 (2021).

3. Georjon, H. & Bernheim, A. The highly diverse antiphage defence systems of bacteria. Nat Rev Microbiol 21, 686–700 (2023).

4. Baca, C.F. & Marraffini, L.A. Nucleic acid recognition during prokaryotic immunity. Molecular Cell 85, 309–322 (2025).

5. Anton, B.P. et al. Biology of host-dependent restriction-modification in prokaryotes. EcoSal Plus, eesp00142022 (2025).

6. Hutinet, G. et al. 7-Deazaguanine modifications protect phage DNA from host restriction systems. Nat Commun 10, 5442 (2019).

7. Crippen, C.S. et al. Deoxyinosine and 7-Deaza-2-Deoxyguanosine as Carriers of Genetic Information in the DNA of Campylobacter Viruses. J Virol 93 (2019).

8. Burke, E.J. et al. Phage-encoded ten-eleven translocation dioxygenase (TET) is active in C5-cytosine hypermodification in DNA. Proc. Natl. Acad. Sci. U. S. A. 118, e2026742118 (2021).

9. Lee, Y.J. & Weigele, P.R. Detection of Modified Bases in Bacteriophage Genomic DNA. Methods Mol Biol 2198, 53–66 (2021).

10. Hutinet, G., Lee, Y.J., de Crecy-Lagard, V. & Weigele, P.R. Hypermodified DNA in Viruses of E. coli and Salmonella. EcoSal Plus 9, eESP00282019 (2021).

11. Rihtman, B. et al. A new family of globally distributed lytic roseophages with unusual deoxythymidine to deoxyuridine substitution. Curr Biol 31, 3199–3206 e4 (2021).

12. Lee, Y.J. et al. Pathways of thymidine hypermodification. Nucleic Acids Res 50, 3001–3017 (2022).

13. Cui, L. et al. Four additional natural 7-deazaguanine derivatives in phages and how to make them. Nucleic Acids Res 51, 9214–9226 (2023).

14. Loenen, W.A. & Raleigh, E.A. The other face of restriction: modification-dependent enzymes. Nucleic Acids Res. 42, 56–69 (2014).

15. Wang, L., Tang, Y., Deng, Z. & Chen, S. DNA Phosphorothioate Modification Systems and Associated Phage Defense Systems. Annu Rev Microbiol 78, 447–462 (2024).

16. Yee, W.X., et al. END nucleases: Antiphage defense systems targeting multiple hypermodified phage genomes. bioRxiv (2025).

17. Monk, I.R., Shah, I.M., Xu, M., Tan, M.W. & Foster, T.J. Transforming the Untransformable: Application of Direct Transformation To Manipulate Genetically Staphylococcus aureus and Staphylococcus epidermidis. MBio 3, e00277 (2012).

18. Xu, S.Y., Corvaglia, A.R., Chan, S.H., Zheng, Y. & Linder, P. A type IV modification-dependent restriction enzyme SauUSI from Staphylococcus aureus subsp. aureus USA300. Nucleic Acids Res. 39, 5597–5610 (2011).

19. Corvaglia, A.R. et al. A type III-like restriction endonuclease functions as a major barrier to horizontal gene transfer in clinical Staphylococcus aureus strains. Proc. Natl. Acad. Sci. U. S. A. 107, 11954–11958 (2010).

20. Ulrich, R.J. et al. Prophage-encoded methyltransferase drives adaptation of community-acquired methicillin-resistant Staphylococcus aureus. J Clin Invest (2025).

21. Bae, T., Baba, T., Hiramatsu, K. & Schneewind, O. Prophages of Newman and their contribution to virulence. Molecular Microbiology 62, 1035–1047 (2006).

22. Tumuluri, V.S., Rajgor, V., Xu, S.Y., Chouhan, O.P. & Saikrishnan, K. Mechanism of DNA cleavage by the endonuclease SauUSI: a major barrier to horizontal gene transfer and antibiotic resistance in Staphylococcus aureus. Nucleic Acids Res. 49, 2161–2178 (2021).

23. Roberts, R.J., Vincze, T., Posfai, J. & Macelis, D. REBASE: a database for DNA restriction and modification: enzymes, genes and genomes. Nucleic Acids Research 51, D629–D630 (2023).

24. Sasnauskas, G., Halford, S.E. & Siksnys, V. How the BfiI restriction enzyme uses one active site to cut two DNA strands. Proc Natl Acad Sci U S A 100, 6410–5 (2003).

25. Lagunavicius, A., Sasnauskas, G., Halford, S.E. & Siksnys, V. The metal-independent type IIs restriction enzyme BfiI is a dimer that binds two DNA sites but has only one catalytic centre. J Mol Biol 326, 1051–64 (2003).

26. Sapranauskas, R. et al. Novel subtype of type IIs restriction enzymes. BfiI endonuclease exhibits similarities to the EDTA-resistant nuclease Nuc of Salmonella typhimurium. J Biol Chem 275, 30878–85 (2000).

27. Sasnauskas, G. et al. A novel mechanism for the scission of double-stranded DNA: BfiI cuts both 3’-5’ and 5’-3’ strands by rotating a single active site. Nucleic Acids Res 38, 2399–410 (2010).

28. Rao, D.N., Dryden, D.T. & Bheemanaik, S. Type III restriction-modification enzymes: a historical perspective. Nucleic Acids Res. 42, 45–55 (2014).

29. Dryden, D.T., Murray, N.E. & Rao, D.N. Nucleoside triphosphate-dependent restriction enzymes. Nucleic Acids Res. 29, 3728–3741 (2001).

30. Loenen, W.A., Dryden, D.T., Raleigh, E.A. & Wilson, G.G. Type I restriction enzymes and their relatives. Nucleic Acids Res. 42, 20–44 (2014).

31. Firman, K. & Szczelkun, M.D. Measuring motion on DNA by the type I restriction endonuclease EcoR124I using triplex displacement. EMBO J 19, 2094–102 (2000).

32. McClelland, S.E., Dryden, D.T. & Szczelkun, M.D. Continuous assays for DNA translocation using fluorescent triplex dissociation: application to type I restriction endonucleases. J Mol Biol 348, 895–915 (2005).

33. Graham, J.E., Sherratt, D.J. & Szczelkun, M.D. Sequence-specific assembly of FtsK hexamers establishes directional translocation on DNA. Proc Natl Acad Sci U S A 107, 20263–8 (2010).

34. Abramson, J. et al. Accurate structure prediction of biomolecular interactions with AlphaFold 3. Nature 630, 493–500 (2024).

35. Liu, L.F. & Wang, J.C. Supercoiling of the DNA template during transcription. Proc Natl Acad Sci U S A 84, 7024–7 (1987).

36. Ostrander, E.A., Benedetti, P. & Wang, J.C. Template supercoiling by a chimera of yeast GAL4 protein and phage T7 RNA polymerase. Science 249, 1261–5 (1990).

37. Szczelkun, M.D., Dillingham, M.S., Janscak, P., Firman, K. & Halford, S.E. Repercussions of DNA tracking by the type IC restriction endonuclease EcoR124I on linear, circular and catenated substrates. EMBO J 15, 6335–47 (1996).

38. Seidel, R. et al. Real-time observation of DNA translocation by the type I restriction modification enzyme EcoR124I. Nat. Struct. Mol. Biol. 11, 838–843 (2004).

39. Torres Montaguth, O.E., Cross, S.J., Lee, L., Diffin, F.M. & Szczelkun, M.D. ENDO-Pore: High-throughput linked-end mapping of single DNA cleavage events using nanopore sequencing. BioRxiv to be submitted (2021).

40. Vitkute, J., Maneliene, Z., Petrusyte, M. & Janulaitis, A. BfiI, a restriction endonuclease from Bacillus firmus S8120, which recognizes the novel non-palindromic sequence 5’-ACTGGG(N)5/4-3’. Nucleic Acids Res. 26, 3348–3349 (1998).

41. Szczelkun, M.D. Kinetic models of translocation, head-on collision, and DNA cleavage by type I restriction endonucleases. Biochemistry 41, 2067–74 (2002).

42. Studier, F.W. & Bandyopadhyay, P.K. Model for how type I restriction enzymes select cleavage sites in DNA. Proc. Natl. Acad. Sci. U. S. A. 85, 4677–4681 (1988).

43. Chand, M.K. et al. Translocation-coupled DNA cleavage by the Type ISP restriction-modification enzymes. Nat Chem Biol 11, 870–7 (2015).

44. Li, Y. et al. Structure and Activation Mechanism of a Lamassu Phage Defence System. bioRxiv, 2025.03.14.643221 (2025).

45. Li, M. et al. Structural insights into type I and type II Lamassu anti-phage systems. bioRxiv, 2025.06.23.660979 (2025).

46. Haudiquet, M. et al. Structural basis for Lamassu-based antiviral immunity and its evolution from DNA repair machinery. bioRxiv, 2025.04.02.646746 (2025).

47. Halford, S.E., Welsh, A.J. & Szczelkun, M.D. Enzyme-mediated DNA looping. Annu Rev Biophys Biomol Struct 33, 1–24 (2004).

48. Dimitriu, T., Szczelkun, M.D. & Westra, E.R. Evolutionary Ecology and Interplay of Prokaryotic Innate and Adaptive Immune Systems. Curr. Biol. 30, R1189–r1202 (2020).

49. Sinkunas, T. et al. Cas3 is a single-stranded DNA nuclease and ATP-dependent helicase in the CRISPR/Cas immune system. EMBO J 30, 1335–42 (2011).

50. Bravo, J.P.K., Aparicio-Maldonado, C., Nobrega, F.L., Brouns, S.J.J. & Taylor, D.W. Structural basis for broad anti-phage immunity by DISARM. Nat Commun 13, 2987 (2022).

51. Huang, P. et al. The mechanism of bacterial defense system DdmDE from Lactobacillus casei. Cell Res 34, 873–876 (2024).

52. Antine, S.P. et al. Structural basis of Gabija anti-phage defence and viral immune evasion. Nature 625, 360–365 (2024).

53. Schwarz, F.W. et al. The helicase-like domains of type III restriction enzymes trigger long-range diffusion along DNA. Science 340, 353–6 (2013).

54. Kulkarni, M., Nirwan, N., van Aelst, K., Szczelkun, M.D. & Saikrishnan, K. Structural insights into DNA sequence recognition by Type ISP restriction-modification enzymes. Nucleic Acids Res 44, 4396–408 (2016).

55. van Aelst, K., Saikrishnan, K. & Szczelkun, M.D. Mapping DNA cleavage by the Type ISP restriction-modification enzymes following long-range communication between DNA sites in different orientations. Nucleic Acids Res 43, 10430–43 (2015).

56. Toliusis, P. et al. The H-subunit of the restriction endonuclease CglI contains a prototype DEAD-Z1 helicase-like motor. Nucleic Acids Res 46, 2560–2572 (2018).

57. Toliusis, P., Zaremba, M., Silanskas, A., Szczelkun, M.D. & Siksnys, V. CgII cleaves DNA using a mechanism distinct from other ATP-dependent restriction endonucleases. Nucleic Acids Res 45, 8435–8447 (2017).

58. Zaremba, M. et al. DNA cleavage by CgII and NgoAVII requires interaction between N- and R-proteins and extensive nucleotide hydrolysis. Nucleic Acids Res 42, 13887–96 (2014).

59. Cohen, H.M., Tawfik, D.S. & Griffiths, A.D. Promiscuous methylation of non-canonical DNA sites by HaeIII methyltransferase. Nucleic Acids Res. 30, 3880–3885 (2002).

60. Ma, B. et al. Biochemical and structural characterization of a DNA N6-adenine methyltransferase from Helicobacter pylori. Oncotarget 7, 40965–40977 (2016).

61. Aranda, J., Roca, M. & Tunon, I. Substrate promiscuity in DNA methyltransferase M. PvuII. A mechanistic insight. Org. Biomol. Chem. 10, 5395–5400 (2012).

62. Borgaro, J.G., Benner, N. & Zhu, Z.Y. Fidelity Index Determination of DNA Methyltransferases. PLoS ONE 8, e63866 (2013).

63. Hall, S.C. & Halford, S.E. Specificity of DNA recognition in the nucleoprotein complex for site-specific recombination by Tn21 resolvase. Nucleic Acids Res 21, 5712–9 (1993).

64. Palmer, B.R. & Marinus, M.G. The dam and dcm strains of Escherichia coli--a review. Gene 143, 1–12 (1994).

65. Yanisch-Perron, C., Vieira, J. & Messing, J. Improved M13 phage cloning vectors and host strains: nucleotide sequences of the M13mp18 and pUC19 vectors. Gene 33, 103–19 (1985).

66. Marko, J.F. & Siggia, E.D. Stretching DNA. Macromolecules 28, 8759–8770 (1995).

67. Lionnet, T., et al. Magnetic trap construction. Cold Spring Harb Protoc 2012, 133–8 (2012).

68. Lionnet, T. et al. Single-molecule studies using magnetic traps. Cold Spring Harb Protoc 2012, 34–49 (2012).

69. Rutkauskas, M., Krivoy, A., Szczelkun, M.D., Rouillon, C. & Seidel, R. Single-Molecule Insight Into Target Recognition by CRISPR-Cas Complexes. Methods Enzymol 582, 239–273 (2017).

70. Chung, S.H. & Kennedy, R.A. Forward-backward non-linear filtering technique for extracting small biological signals from noise. J Neurosci Methods 40, 71–86 (1991).

