## Supplementary Figures for "Individual *Staphylococcus aureus* SauUSI restriction endonuclease motors fragment methylated DNA"

Szczelkun, M.D.

**Supplementary Fig. 1| Effect of different traps on slowing reinitiation of translocation following dissociation during initiation.**

**Supplementary Fig. 2| Fitting the triplex displacement data to an ordinary differential equation model is consistent with upstream initiation.**

**Supplementary Fig. 3|A rare codirectional translocation event.**

**Supplementary Fig. 4|Temperature-dependence of the translocation rate.**

**Supplementary Fig. 5|SauUSI can cleave DNA lacking free termini.**

**Supplementary Fig. 6|DNA ends generated by primary cleavage are similar for all conditions.**

**Supplementary Fig. 7|Strand-dependent cleavage distributions.**

**Supplementary Fig. 8|Dinucleotide preferences at the distal cleavage site.**

**Supplementary Fig. 9| SauUSI cleavage distributions are translocation strand-dependent.**

**Supplementary Fig. 10| dCas9 provides an orientation-dependent roadblock to SauUSI translocation.**

**Supplementary Fig. 11|DNA translocation and cleavage are confined *in cis* to methylated DNA.**

**Supplementary Fig. 12|Calibration of the phosphate binding protein sensor.**

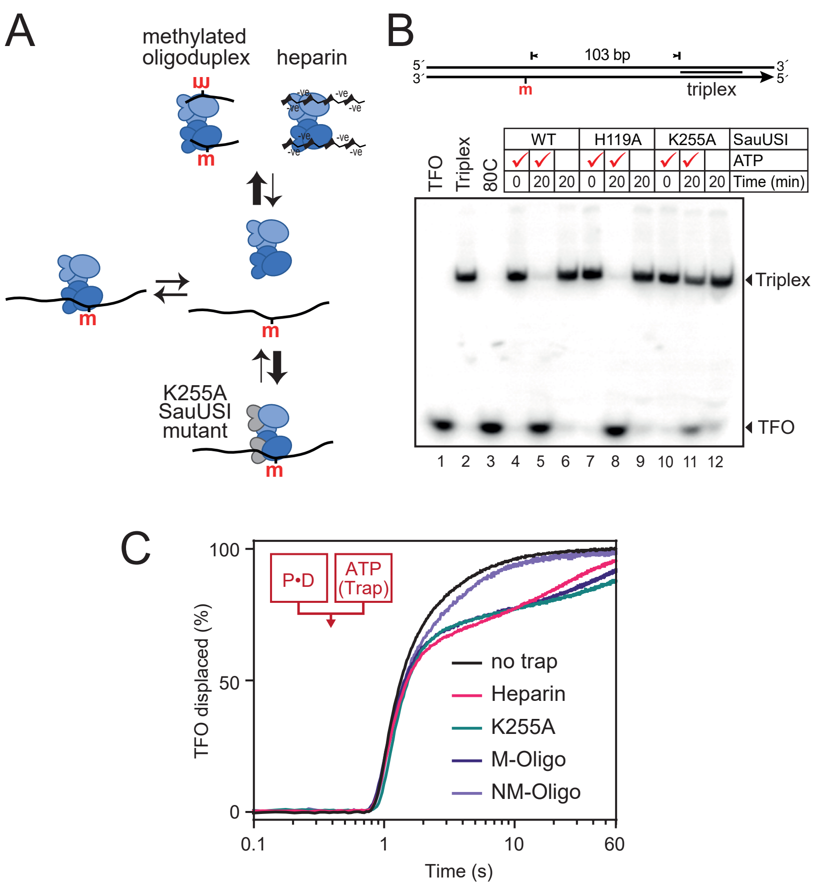


**Supplementary Fig. 1| Effect of different traps on slowing reinitiation of translocation following dissociation during initiation. A**, Model for inhibition of reassociation of SauUSI and DNA following dissociation using traps that sequester free SauUSI (M-Oligo or heparin) or free DNA (K255A Walker A mutant of SauUSI). **B**, Displacement of a triplex forming oligonucleotide (TFO) by different SauUSI proteins using a triplex gel electrophoresis assay with hemimethylated DNA. The nuclease mutant H119A can fully displace the triplex while the ATPase mutant K255A produces only partial displacement, indicating a low background of translocation activity, making it suitable as a trap. **C**, Time-dependent triplex displacement from hemimethylated linear DNA (103 bp spacing, panel B) using a stopped flow assay. Samples were mixed as indicated (cartoon) with Protein (P), DNA (D), ATP and 5 µM Heparin, 1 µM oligoduplex or 0.25 µM K255A helicase mutant dimer, where included. M-Oligo is the hemimethylated oligoduplex, NM-Oligo is a non-methylated oligoduplex.

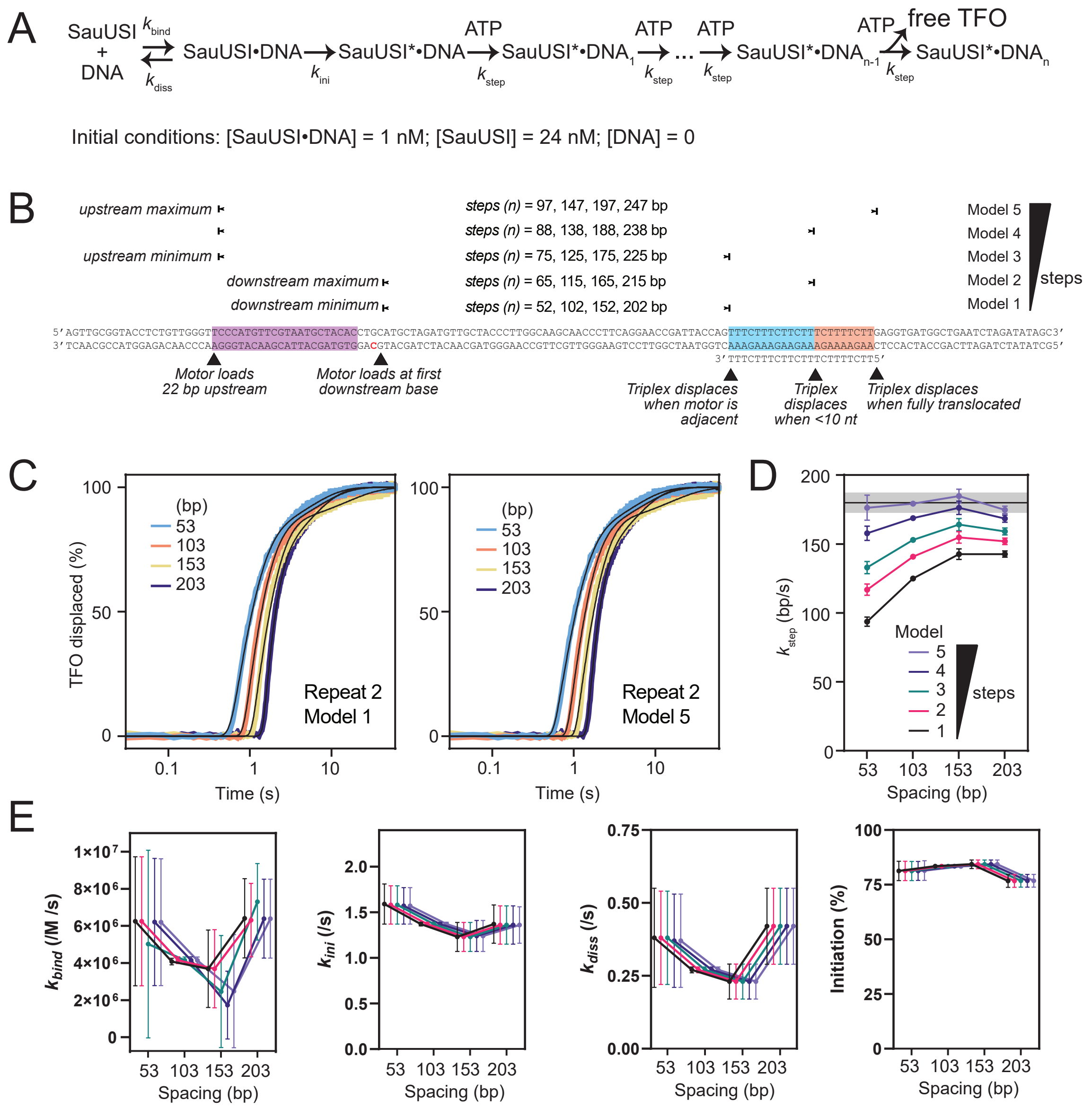


**Supplementary Fig. 2| Fitting the triplex displacement data to an ordinary differential equation model is consistent with upstream initiation. A**, Model for triplex displacement described that was sufficient to describe the data. Dissociation during translocation was considered too infrequent to consider. Initial conditions were chosen to match the triplex assays in **Fig. 1C**. **B**, Based on the substrates used in **Fig. 1C,** five models were tested with different initiation sites representing down- or up-stream initiation and with different numbers of steps required for triplex displacement. **C**, Models were coded as ODEs in Berkely Madonna and the relevant number of steps fitted to each data repeat, with the $k_{step}$, $k_{ini}$, $k_{bind}$ and $k_{diss}$ parameters floated. Examples are shown for the fits between the models with the fewest (1) and most (5) number of steps and one repeat of the four triplex spacings. **D**, Mean $k_{step}$ values (3 repeats, error bars SD) vary less as the number of steps increases and only model 5 approaches the average value determined from the lag phases in **Extended Data Fig. 1** (horizontal line is 180 bp/s, grey box represents the SD error of ±7 bp/s). **E**, Other parameters are independent of the number of steps but vary more with data error.

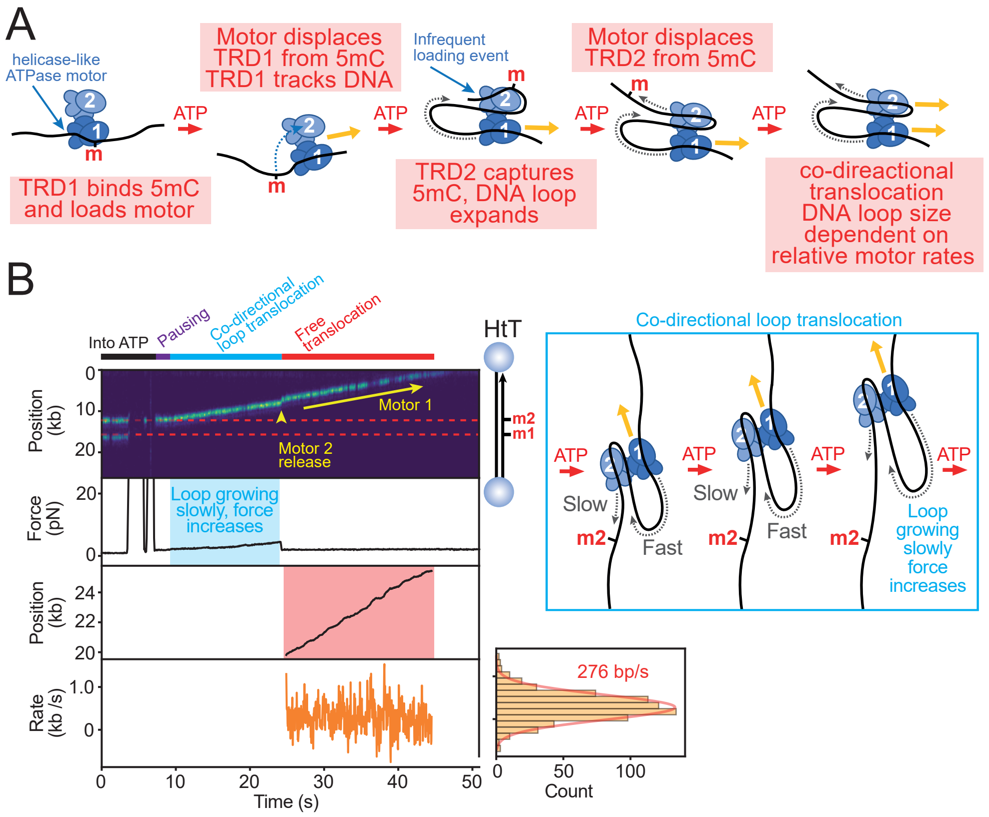


**Supplementary Fig. 3|A rare codirectional translocation event. A,** Model for how initiation at a hemimethylated site can lead to the loading of two motors that translocate in the same direction. Key to this is that rather than when TRD2 captures the displaced 5mC, the DNA can occasionally bind in a configuration that allows the second helicase motor to load. This results in both motors translocating 3′-5′ on the methylated strand. **B**, Representative kymograph at 1.5 pN showing a dual hemimethylated HtT DNA-bead tether with labelled enzymes (655 nm quantum dot-streptavidin-biotin-WT SauUSI) prebound at each 5mC. Following a brief delay after moving into the ATP channel, one SauUSI dissociates while the second initiates loop translocation with a corresponding increase in force that is lower than expected for unidirectional loop translocation based on the observed rate of motion. Where two co-directional motors translocate at the same rate, the force would not be expected to increase as the loop size is not changing. The slowly increasing force can be rationalized as the leading motor having a moderately higher rate than the following motor, resulting in a loop that grows slowly (inset cartoon). Release of motor 2 produces a jump “forward” (release of motor 1 would have produced a jump “backward”) and results in free translocation at the expected rate (**Extended data Fig. 1A**).

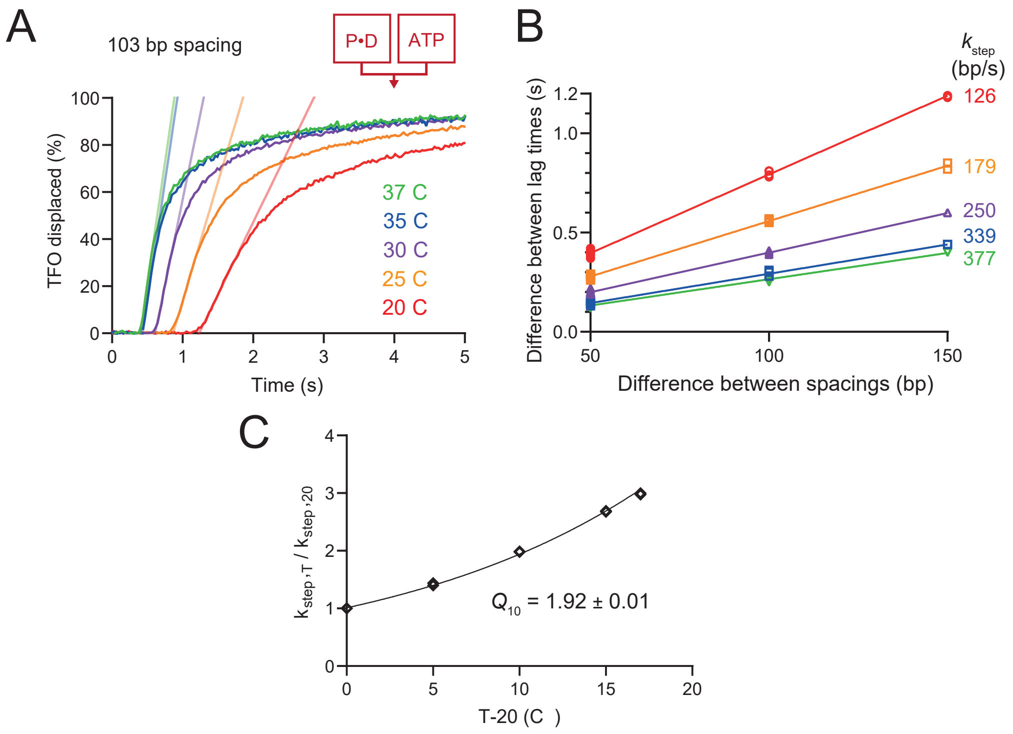


**Supplementary Fig. 4|Temperature-dependence of the translocation rate**. **A**, Representative time-course of triplex displacement with a 103 bp 5mC-triplex spacing at different temperatures measured using stopped flow. Diagonals are linear fits to estimate the lag time. **B**, Using DNA in **Fig. 1C** to obtain lag times at different temperatures, rates were estimated from linear fits. **C**, Estimation of the Q10 temperature coefficient (Mean and standard deviation) from two data repeats.

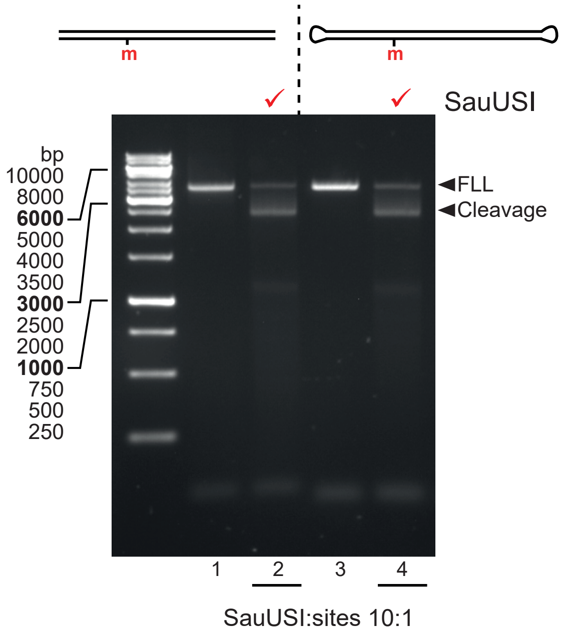


**Supplementary Fig. 5|SauUSI can cleave DNA lacking free termini.** Linear DNA with free or hairpins ends was cleaved with excess SauUSI for 10 min and analyzed by agarose gel electrophoresis. Both DNA were cleaved to the same extent.

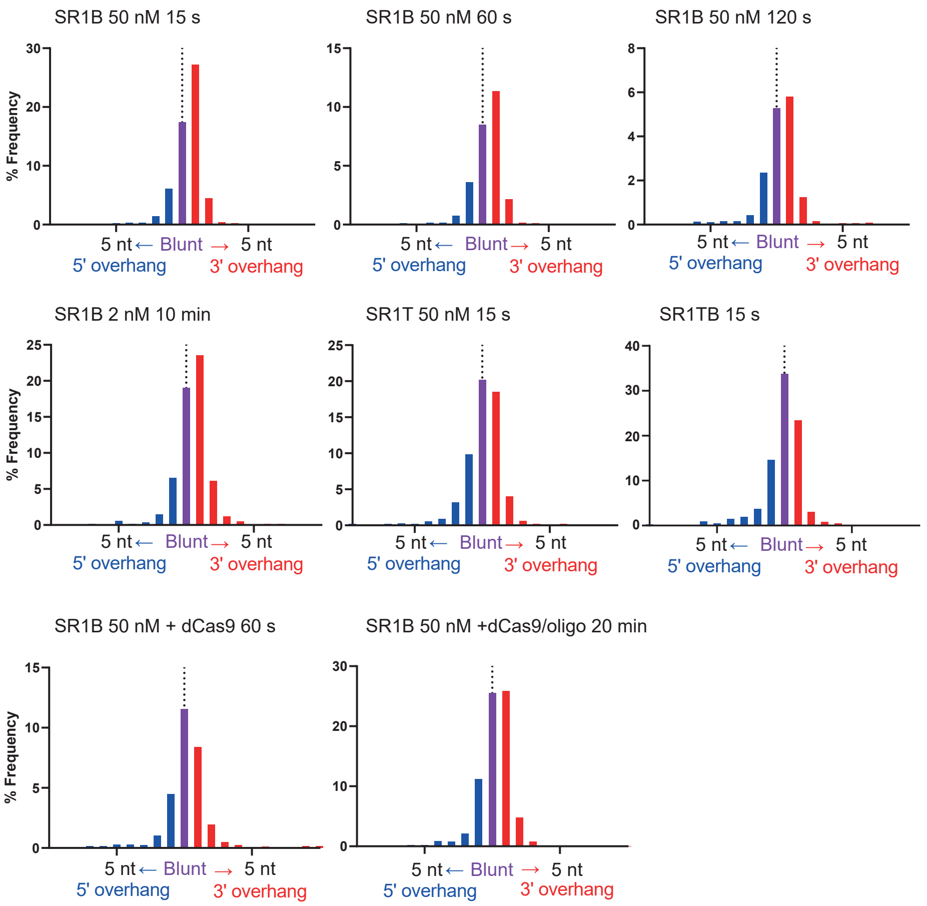


**Supplementary Fig. 6|DNA ends generated by primary cleavage are similar for all conditions.** Comparison of the distribution of primary cleavage site spacings from ENDO-Pore experiments with the DNA and conditions indicated (1 nt bins). Differences in relative amounts of blunt and 1 nt 3' extensions reflects sampling of different strands or parts of the substrate rings with different experimental conditions and/or substrates.

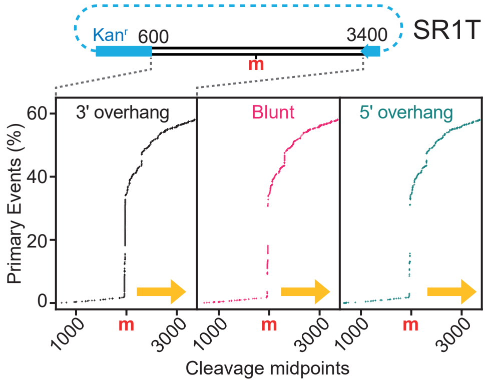


**Supplementary Fig. 7 | Strand-dependent cleavage distributions.** Cumulative percentage of primary 3′ extension, blunt or 5′ extension events relative to the 5mC position (m) on SR1B. Cleavage within the Kanamycin resistance cassette cannot be mapped by ENDO-Pore. Yellows arrow indicates polarity of unidirectional translocation. While some proximal loci favor one sort of end, reflecting preferences for particular top/bottom strand positions, distal cleavages are equally distributed.

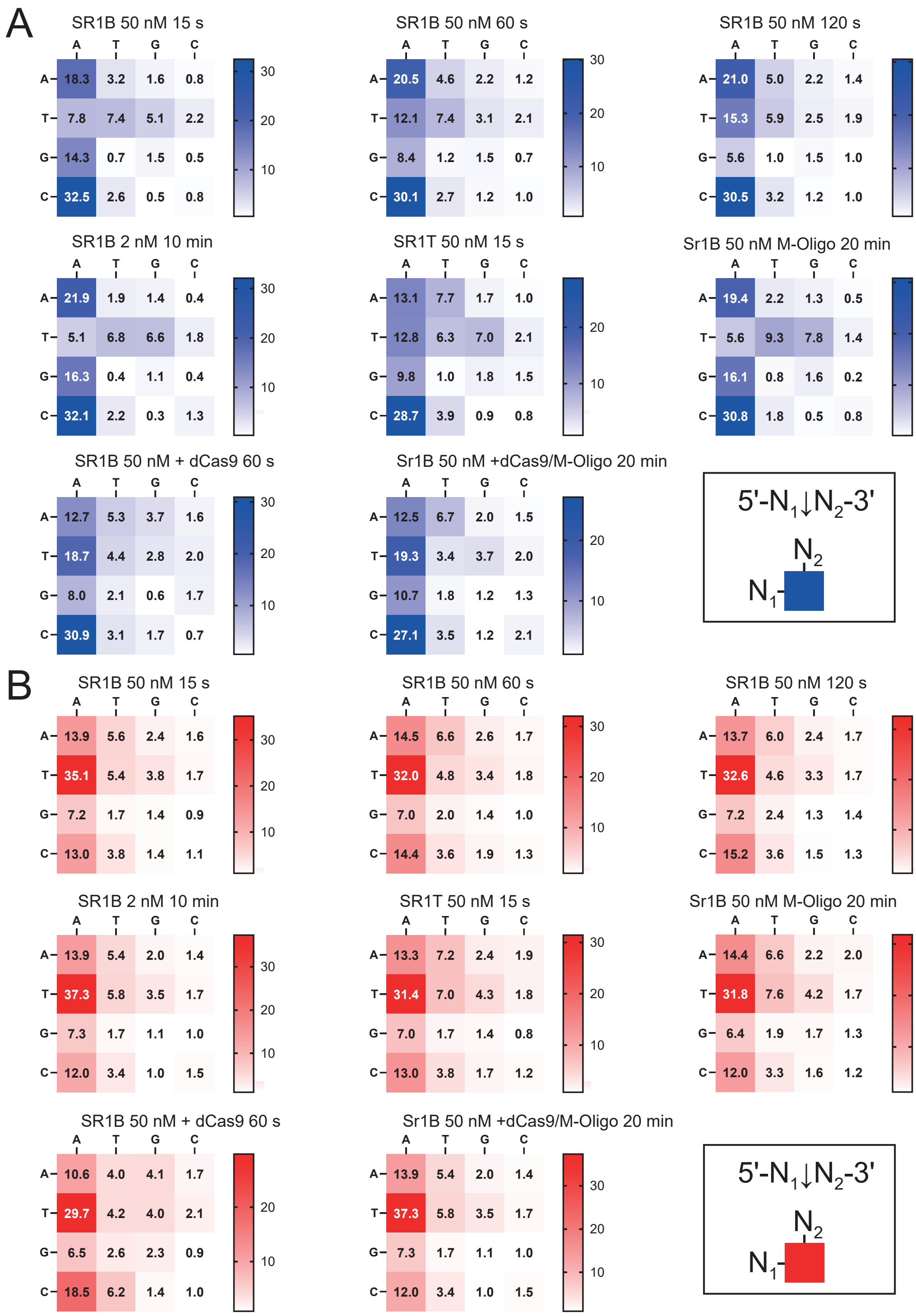


**Supplementary Fig. 8|Dinucleotide preferences at the distal cleavage site**. Comparison of the percentage of dinucleotide distributions at distal primary and secondary DNA cleavage sites from ENDO-Pore experiments, with the DNA and conditions indicated. For the primary cleavages, there are differences observed where cleavage loci are limited by the dCas9 roadblock or where translocation is on the opposite strand (SR1T) that likely reflect the different sites that the nuclease can access. For the secondary distal cleavages, the distributions are more similar, and more strongly favors a 5′-TA-3′, suggesting that these sites are cleaved more slowly than the 5′-CA-3′ > 5′-AA-3′ > 5′-GA-3′ sites that are preferred during earlier primary cleavage.

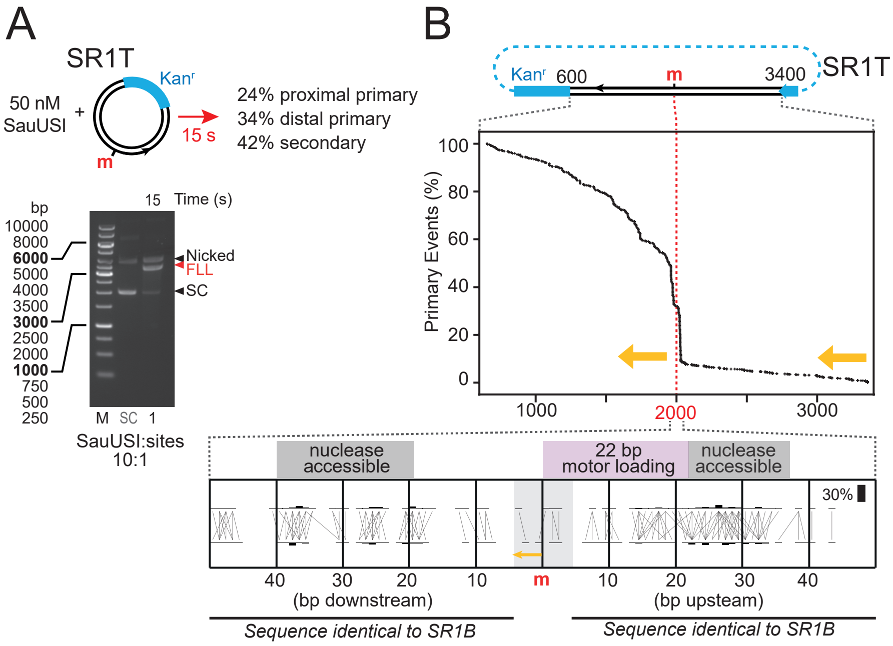


**Supplementary Fig. 9 | SauUSI cleavage distributions are translocation strand-dependent. A**, Cartoon shows SR1T map. Agarose gel of 15 s reaction used for ENDO-Pore. Event type percentages are quantified from the ENDO-Pore data. **B**, Cumulative percentage of primary events on SR1T relative to the 5mC position (m) – note the opposite polarity of translocation (yellow arrow). For the proximal region, a strand linkage plot (SLP) shows bars representing the percentage of phosphodiester cleavage at each top and bottom location and horizontal/vertical lines that indicate the ends generated (blunt or overhangs). The up- and downstream sequences are the same as SR1B except for the grey box. Note, however, that the motor is translocating on the opposite strand and this may partly explain the different relative cleavages compared to SR1B.

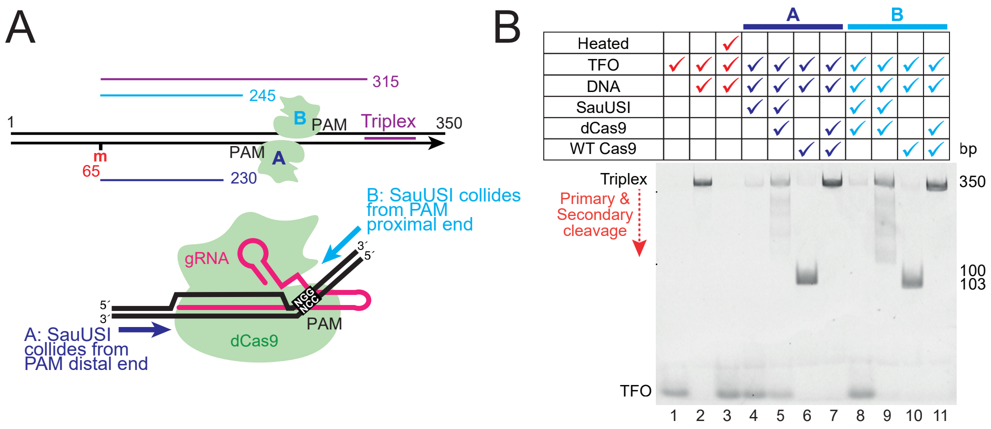


**Supplementary Fig. 10| dCas9 provides an orientation-dependent roadblock to SauUSI translocation. A**, Schematic of triplex DNA used to test if dCas9 blocked SauUSI translocation. Two separate gRNAs were used, gRNA-A and gRNA-B, which are at similar distances from the methylated site but PAMs on opposite strands, flipping the orientation of the dCas9:gRNA:DNA complex (i.e. PAM proximal or PAM distal), highlighted in the cartoon (lower panel). **B**, A triplex displacement gel to determine whether SauUSI can bypass dCas9. DNA shredding is observed in lanes 5/9, which is predicted to be due to dCas9 blocking translocation past the dCas9 complex and capturing cleavage events upstream of the dCas9 placements. Displacement is reduced or completely absent in lanes 5 and 9, respectively. This suggests that PAM proximal gRNA-B could effectively block SauUSI translocation whereas PAM distal gRNA-A only partially blocked SauUSI translocation.

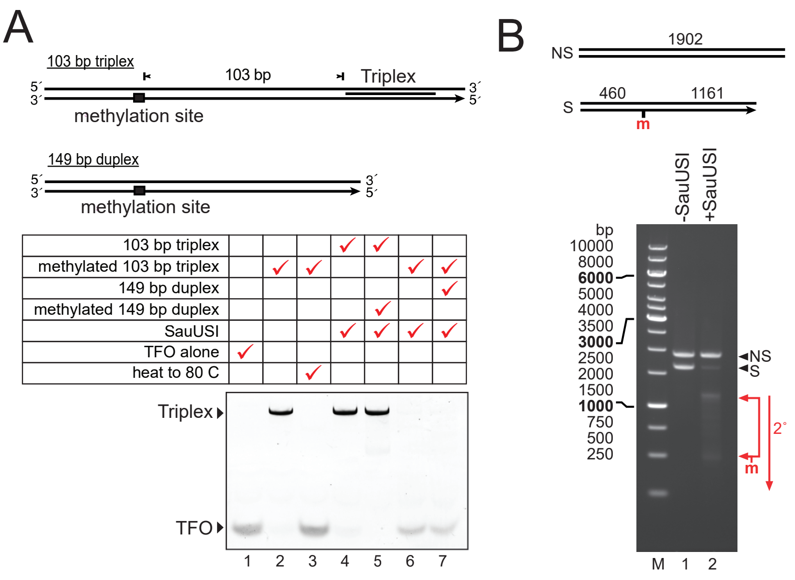


**Supplementary Fig. 11|DNA translocation and cleavage are confined *in cis* to methylated DNA. A**, Triplex displacement activity cannot transfer from methylated to non-methylated DNA. Displacement of a TFO was measured using a triplex gel electrophoresis assay with methylated or non-methylated DNA (M1.BfuAI) in the presence of methylated or non-methylated DNA without a triplex bound. **B**, Primary and Secondary DNA cleavage is confined to the methylated DNA. 20-minute cleavage by excess SauUSI of a hemimethylated substrate in the presence of an unmethylated DNA, measured using agarose gel electrophoresis.

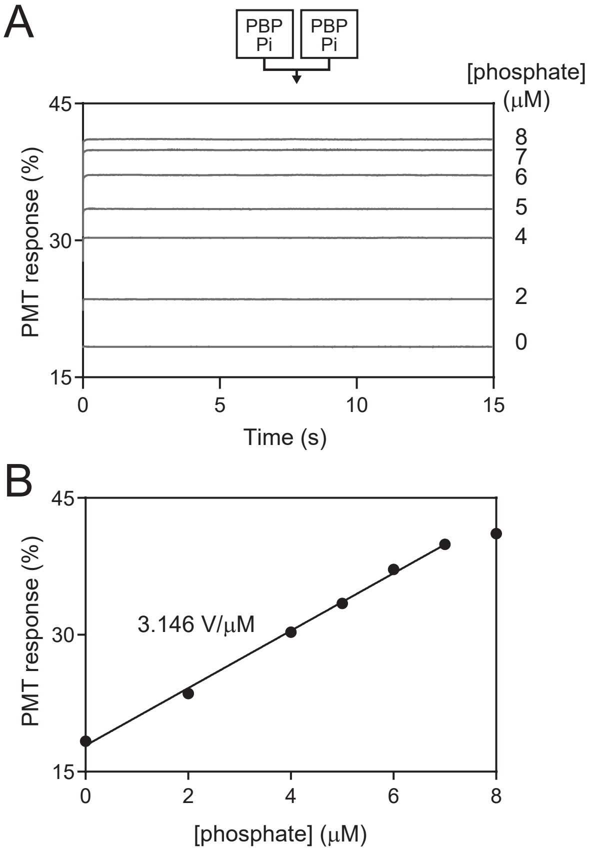


**Supplementary Fig. 12|Calibration of the phosphate binding protein sensor.** A, Raw fluorescence signal intensity (in photomultiplier volts) in the stopped flow at different phosphate concentrations. Samples were mixed as indicated (cartoon) with both syringes containing 6 µM phosphate binding protein (PBP) and phosphate (Pi). **B**, Calibration plot of the average signals from Panel A at fixed concentrations of phosphate. The slope of the linear regression was used to convert the observed signal to released phosphate and hence ATPs hydrolyzed.
