## Supplementary Methods for "Individual *Staphylococcus aureus* SauUSI restriction endonuclease motors fragment methylated DNA"

**Protein Purification Methods**

See Supplementary Excel file for synthetic gene and oligonucleotide sequences

*SauUSI variants*

A codon optimised wild-type SauUSI gene was acquired from Integrated DNA Technologies in pUCIDT. The gene was subcloned using BsaI and BamHI into the BsaI and BamHI-treated pE-SUMO Amp (LifeSensors). pE-Avi SauUSI was generated by replacing the His-SUMO-tag with the Avi-Tag (Avidity LLC) using overlap extension PCR. Mutants were generated by site-directed mutagenesis using overlapping primers.

Wild-type and mutant SUMO-tagged SauUSI proteins were purified using the same protocol. *E. coli* BL21 (DE3) cells (ThermoFisher) were transformed with the vector and single colonies selected on LB agar with 50 µg/ml ampicillin. Two 2.5-litre flasks of 500 mL LB with 50 µg/ml ampicillin were inoculated with 5 mL of an overnight starter culture and grown at 37 °C and 200 rpm until $OD_{600nm}$ of 0.6. IPTG was added to 1 mM and cells incubated for 18-20 h at 18 °C. Overnight cultures were pelleted at 6238*g* for 30 minutes at 4 °C, washed with 25 mL of 10 mM Tris-Cl (pH 8.0), 100 mM NaCl and 1 mM ethylenediaminetetraacetic acid (EDTA) per flask, and cells re-pelleted at 4,500*g* for 20 minutes at 4 °C.

Cell pellets were resuspended in 10 mL/g of 50 mM Tris-Cl (pH 8.0), 500 mM NaCl, 30 mM Imidazole, 5 mM $MgCl_2$, 10% (v/v) Glycerol, 1 mM β-mercaptoethanol (β-ME) and EDTA-free protease inhibitor tablets (Roche). Cells were sonicated using a Sonics Vibra-Cell sonicator fitted with a 0.5-inch tip at 65% amplitude, with 2 s on and 2 s off for 2 minutes and cleared at 6238*g* for 30 minutes at 4 °C. Clarified lysates were dialyzed against 2 L of Buffer-HisA (50 mM Tris-Cl (pH 8.0), 500 mM NaCl, 30 mM Imidazole and 1 mM β-ME) using 10-kDa molecular weight cut-off (MWCO) SnakeSkin dialysis tubing (Pierce) for 2 h at 4 °C. The dialysate was loaded onto a 5-mL His-TALON cobalt metal affinity column (Takara Bio) pre-equilibrated with buffer His A. SauUSI was eluted using a step gradient of His A supplemented with a final concentration of 140 mM Imidazole at 4 mL/min and fractions selected by SDS-polyacrylamide gel electrophoresis. Pooled fractions were dialyzed against 2 litres of Buffer-His B (50 mM Tris-Cl (pH 8.0), 200 mM NaCl and 1 mM β-ME) for >16 h at 4 °C and simultaneously digested with SUMO protease to remove the His-SUMO tag.

The supernatant was loaded onto a 5-mL His-TALON column pre-equilibrated with buffer His A. The flow-through was collected and diluted 1:1 with buffer MQ0 (20 mM Tris-Cl (pH 8.0), 0.1 mM EDTA and 1 mM β-ME) before loading onto an 8-mL MonoQ 10/100 GL column (Cytiva) pre-equilibrated with MQ1 (MQ0 supplemented with 150 mM NaCl) at 4 mL/min. SauUSI eluted as two peaks over a linear gradient from 150 mM to 500 mM NaCl at 4 mL/min. The second peak (dimeric

SauUSI) was selected using SDS-polyacrylamide gel electrophoresis and concentrated to <1 mL using an Amicon Ultra-15 centrifugal filter with a 100 kDa MWCO at 3,500*g* and 4 °C before storage as a 50% (v/v) glycerol stock at -20 °C.

Biotinylated SauUSI variants were expressed as described for the His-SUMO SauUSI, except: cells were co-transformed with pBirA (Avidity) and single colonies isolated on LB agar with 50 µg/mL ampicillin and 34 µg/mL chloramphenicol; and, cultures were induced as above plus the addition of 50 µM Biotin. The same protocol was used for harvesting and sonication except for the buffer (100 mM Tris-Cl (pH 8.0), 500 mM NaCl, 5 mM $MgCl_2$, 2.5 mM EDTA, 1 mM DTT and EDTA-free protease inhibitor tablet). After clarification, the supernatant was dialysed against buffer SLA (10 mM Tris (pH 8.0), 150 mM NaCl, 0.1 mM EDTA & 1 mM DTT) for 2 h at 4 °C. The dialysate was loaded onto a pre-equilibrated 10 mL Softlink Avidin column (Promega) at 1 mL/min. Following a wash of buffer SLB (SLA with the addition of 5 mM biotin) for 5 minutes at 1 mL/min, the column was incubated without flow for 30 minutes at 4 °C before protein elution at 1 mL/min using buffer SLB. Fractions were selected using SDS-polyacrylamide gel electrophoresis and dialysed into MQ1 for >2 h at 4 °C. The dialysate was loaded onto a pre-equilibrated 8 mL MonoQ 10/100 column and eluted as above; the remaining steps were the same as the SUMO variant.

*AluI Methyltransferase*

A synthetic, codon optimised AluI methyltransferase (M.AluI) gene was obtained from Twist Biosciences and subcloned into pE-SUMO using PCR amplification and BsaI Golden Gate cloning (New England Biolabs). T7 express *E. coli* cells (New England Biolabs) were transformed with pE-SUMO M.AluI and single colonies isolated on LB agar with 50 µg/mL ampicillin. Two 2.5 Litre flasks containing 500 mL LB with 50 µg/mL ampicillin were inoculated and grown at 37 °C until $OD_{600nm}$ of 0.6. Following addition of 1 mM IPTG, cultures were grown at 18 C for 18-20 h before harvesting as described above. The purification scheme and buffers are consistent with the His-SUMO SauUSI protocol previously described, except only one M.AluI peak was observed during the MonoQ step.

*M1.BfuAI and M2.BfuAI Methyltransferases*

Synthetic codon optimised genes for wild-type BfuAI methyltransferases 1 (M1.BfuAI) and 2 (M2.BfuAI) were obtained from Integrated DNA Technologies. M1.BfuAI was subcloned by treating with BsaI and HindIII followed by ligation into BsaI and HindIII-treated pE-SUMO. M2.BfuAI was subcloned by treating with BsaI and XbaI followed by ligation into BsaI and XbaI-treated pE-SUMO. The SUMO-tag was removed by overlap extension PCR to generate pE-His-M2.BfuAI. T7 express *E. coli* cells were transformed with either plasmid, and single colonies isolated on LB agar with 50 µg/mL ampicillin. For both proteins, two 2.5 litre flasks containing 500 mL LB with 50 µg/mL ampicillin were inoculated with

5 mL of an overnight starter culture and grown at 37 °C. For M1.BfuAI, cells were grown until $OD_{600nm}$ of 0.8 and induced with 1 mM IPTG. Following induction, cells were incubated at 37 °C for 2 h before being harvested as above. For M2.BfuAI, cells were grown until $OD_{600nm}$ of 0.6 and induced with 1 mM IPTG. Following induction, cells were incubated at 18 C for 18-20 h before being harvested as above.

His-SUMO M1.BfuAI cells were resuspended in 10 ml/g of 50 mM HEPES (pH 7.0), 500 mM NaCl, 5 mM $MgCl_2$, 0.5 mM EDTA, 1 mM β-ME and EDTA-free protease inhibitor tablet (Roche). Cells were sonicated and clarified using the same protocol described for SauUSI. The clarified supernatant was dialysed against buffer His C (20 mM HEPES (pH 7.0), 300 mM NaCl, 5 mM Imidazole and 1 mM β-ME) for 2 h at 4 °C. The dialysate was loaded onto a 5-mL His-TALON column pre-equilibrated with buffer His C at 4 ml /min before elution with His C supplemented with a final concentration of 130 mM Imidazole. Fractions were selected using SDS-polyacrylamide gel electrophoresis before dialysis for >16 h at 4 °C into buffer His C, with the addition of SUMO protease to remove the SUMO tag.

The dialysed and cleaved M1.BfuAI was loaded onto a 5 mL His-TALON column pre-equilibrated with buffer His C at 4 mL/min and the flowthrough collected and dialysed in buffer MQ2 (20 mM HEPES (pH 7.0), 100 mM NaCl, 0.1 mM EDTA and 1 mM DTT) for 2 h at 4 °C. The dialysate was loaded onto a MonoQ 10/100 GL column pre-equilibrated with buffer MQ2 at 4 mL/min before elution with a linear gradient between 100 mM to 550 mM NaCl. Fractions containing M1.BfuAI were determined by SDS-polyacrylamide gel electrophoresis and concentrated to <1 mL using an Amicon Ultra-15 centrifugal filter with a 10 kDa MWCO at 3,500*g* and 4 °C, before storage as a 50% (v/v) glycerol stock at -20 °C.

His-M2.BfuAI cells were resuspended in 10 mL/g of 50 mM Tris (pH 8.0), 500 mM NaCl, 5 mM $MgCl_2$, 0.5 mM EDTA, 1 mM β-ME and EDTA-free protease inhibitor tablet (Roche). Cells were sonicated and clarified using the same protocol described for SauUSI. The clarified supernatant was dialysed against buffer His D (50 mM Tris (pH 7.4), 300 mM NaCl, 5 mM Imidazole and 1 mM β-ME) for 2 h at 4 °C. After dialysis, dialysate was loaded onto a 5 mL His-TALON column pre-equilibrated with buffer His D at 4 mL/min followed by elution with His D supplemented with a final concentration of 125 mM Imidazole. Fractions were identified using SDS-polyacrylamide gel electrophoresis and dialysed against buffer MS1 (50 mM Tris (pH 7.4), 125 mM NaCl, 0.1 mM EDTA and 1 mM DTT) for >16 h at 4 °C. The dialysate was loaded onto a 1 mL MonoS 5/50 column (Cytiva) pre-equilibrated with buffer MS1 at 4 mL/min before elution with a linear gradient between 125 mM to 563 mM NaCl. Fractions containing His-M2.BfuAI were selected by SDS-polyacrylamide gel electrophoresis and concentrated and stored as described for M1.BfuAI.

*Tn21 resolvase*

A synthetic codon optimised gene for Tn*21* resolvase was obtained from Integrated DNA Technologies and was subcloned by treating with BsaI and BamHI followed by ligation into BsaI and BamHI-treated pE-SUMO. BL21 (DE3) STAR cells (ThermoFisher) were transformed with pE-His-Sumo-Tn21 and single colonies isolated on LB agar with 100 µg/mL ampicillin. Two 2.5 litre flasks containing 500 mL LB with 100 µg/mL ampicillin were inoculated with 5 mL of an overnight starter culture and grown at 37 °C until $OD_{600nm}$ 0.4 – 0.6. IPTG was added to 1 mM and cells incubated for 18-20 h before being harvested as above.

Cell pellets were resuspended in 10 ml per gram of 50mM Tris-Cl (pH7.4), 500 mM NaCl, 0.5 mM EDTA and EDTA-free protease inhibitor tablet (Roche). Cells were sonicated and clarified as above before dialysis against Buffer A (50 mM Tris (pH 7.4), 300 mM NaCl and 5 mM Imidazole). Dialysate was loaded onto a 5 mL His-TALON column pre-equilibrated with Buffer A at 5 mL/min followed by elution with Buffer A supplemented with a final concentration of 300 mM Imidazole at 1 mL/min. Fractions were identified using SDS-polyacrylamide gel electrophoresis and dialysed with SUMO-protease against Buffer C (50 mM Tris (pH 7.4), 500 mM NaCl) for >16 h at 4 °C.

The dialysed and cleaved Tn*21* resolvase was diluted to 400 mM NaCl and supplemented with 0.1 mM EDTA and 10% (v/v) glycerol before loading onto a 1 mL MonoQ 5/50 column pre-equilibrated with 50 mM Tris (pH 7.4), 400 mM NaCl, 0.1 mM EDTA and 10% (v/v) glycerol at 1 mL/min. Protein was eluted with a linear gradient to 1 M NaCl and Tn*21* fractions identified by SDS-polyacrylamide gel electrophoresis. Pooled Tn*21* was diluted to 500 mM NaCl and supplemented with 1 mM final EDTA followed by slow addition of 0.2 g/mL ammonium sulphate with gentle mixing. The supernatant was clarified at 3000*g* for 30 min at 4 °C and a further 0.2 g/mL ammonium sulphate was slowly added with gentle mixing. Precipitated protein was collected at 3000*g* for 30 min at 4 °C. The pellet was resuspended and stored in 50 mM Tris (pH 7.4), 500 mM NaCl, 0.1mM EDTA, 50% (v/v) glycerol.

*Phosphate Binding Protein*

Fluorescently labelled phosphate binding protein (MDCC-PBP) was prepared and purified as described previously[53].

*Converting force to loop size*

During loop translocation, the apparent contour length is reduced by approximately the size of the expanding loop. At a fixed trap-to-trap distance and constant power, this transiently reduces the end-to-end extension, increasing DNA tension and thus observed force. To use measured force changes and end-to-end extension to back-calculate changes in the apparent contour length, and thus loop size, we used the Marko–Siggia interpolation formula for the Worm-Like Chain (WLC) model for DNA elasticity that is reliable at low forces[66]:

$$\frac{FA}{k_BT} = \frac{1}{4\left(1 - \frac{Z}{L}\right)^2} - \frac{1}{4} + \frac{Z}{L}$$

Where *F* is the force, *A* is the DNA persistence length (51.35 nm), $k_B$ is the Boltzmann constant (1.38e-23 J/K), T is temperature in Kelvin (ambient at 295 K), *Z* is the stretched end-to-end DNA extension and *L* is the contour length of the DNA. Solving for *L* using https://www.wolframalpha.com/ gives:

$$L = \frac{\left(54X^3Z^3 - 27X^2FZ^3 + 3\sqrt{3}\sqrt{135X^4F^2Z^6 - 108X^3F^3Z^6 + 144X^2F^4Z^6 - 64XF^5Z^6} + 72XF^2Z^3 - 16F^3Z^3\right)^{1/3}}{6 \cdot 2^{1/3}F} - \frac{-36X^2Z^2 + 12XFZ^2 - 16F^2Z^2}{12 \cdot 2^{2/3}F\left(54X^3Z^3 - 27X^2FZ^3 + 3\sqrt{3}\sqrt{135X^4F^2Z^6 - 108X^3F^3Z^6 + 144X^2F^4Z^6 - 64XF^5Z^6} + 72XF^2Z^3 - 16F^3Z^3\right)^{1/3}} + \frac{3XZ + 4FZ}{6F}$$

where:

$$X = \frac{A}{k_BT}$$

This returns real and imaginary parts where the latter is extremely small (essentially zero in practical terms). We implemented this equation in the ctraptools package (https://github.com/sjcross/ctraptools).
